# Autophagy suppression by TORC1 maintains epithelial plasma membrane integrity and inhibits syncytium formation

**DOI:** 10.1101/2021.08.05.455228

**Authors:** Parisa Kakanj, Sourabh Bhide, Bernard Moussian, Maria Leptin

## Abstract

Epithelial wound healing in *Drosophila* involves the formation of multinucleate cells surrounding the wound. We show that autophagy, a cellular degradation process often deployed in stress responses, is required for the formation of a multinucleated syncytium during wound healing. In addition, uncontrolled autophagy in the unwounded epidermis leads to the degradation of endo-membranes and the lateral plasma membrane, while the apical and basal membranes and the epithelial barrier function remain intact. Proper functioning of TORC1 is needed to prevent autophagy from destroying the larval epidermis, which depends on membrane isolation and phagophore expansion, but does not require the fusion of autophagosomes to lysosomes. Our findings reveal a function for TORC1-mediated regulation of autophagy in maintaining membrane integrity and homeostasis in the epidermis and during wound healing. Finally, autophagy can counteract experimentally induced nuclear defects resembling laminopathies.

**Key findings:**

1. A novel role for TORC1/autophagy pathway to control plasma membrane integrity and homeostasis.
2. Autophagy as the only known necessary and sufficient inducer of syncytium formation in the epithelium and during wound healing.

## Introduction

Autophagy is a conserved intracellular degradation process that engulfs cytoplasmic materials in double membrane vesicles, the autophagosomes, and delivers them to lysosomes^1^. Multiple organelles such as the endoplasmic reticulum (ER), Golgi, mitochondria, nuclear envelope and plasma membrane can provide membrane to form pre-autophagosomal structures (PAS) and nucleate phagophores^2, 3^. Autophagy occurs during development, differentiation and tissue remodelling, and it can be activated in response to intracellular or extracellular stresses such as DNA damage, protein aggregates, damaged organelles, starvation, heat shock, oxidative stress infection or wounding^4, 5^.

Wounding has been shown in *Drosophila*, planarians, zebrafish, rat, mouse and plants to induce autophagy^1, 6–12^. In plants, rats and mice, autophagy is in turn necessary for efficient wound healing. Plant grafting induces autophagy at the site of the wound and lack of Atg2 or Atg5 reduces the rate of successful grafting^7^. Wounding mouse skin induces the expression of autophagy genes in the epidermis and some of these genes (Atg5 or Atg7) are needed for proper wound closure, infiltration by immune cells, and for the production of cytokine CCL2. Applying recombinant CCL2 can reverse the effect of impaired autophagy indicating that the main role of autophagy in this context is to promote wound healing by inducing the production of CCL2^11^. In rats, elevated autophagy at the wound margin is required for clearance of bacteria. Both autophagy and bacterial clearance are reduced in diabetic rats, but can be ameliorated by inhibiting TOR signalling with rapamycin^12^. These findings illustrate a role for autophagy in wound healing, but many of the molecular mechanisms involved are unclear. Here, we used *Drosophila* larvae to investigate the cellular function and physiological role of autophagy in the epidermis and during wound healing.

The epidermis of *Drosophila* larva is a monolayer of polyploid, postmitotic epithelial cells on a basal lamina, attached on its apical side to the cuticle^13, 14^. Larval wound healing is driven by two parallel and coordinated mechanisms. Actomyosin assembly and contraction at the apical side and lamellipodia on the basal side of the cells surrounding the wound lead to elongation of these cells into the wound to close the gap^6, 13^. Autophagy occurs in cells surrounding the wound as they close the wound^6^. Another late event in wound healing in larvae, pupa and adult flies is the formation of a multinucleate syncytium^13, 15, 16^.

One of the pathways activated during wound healing is the JNK pathway. Hyperactivation of JNK leads to disassembly of focal adhesions complexes and induces syncytium formation in unwounded epidermis^13, 15^. However, the natural syncytium formation in cells surrounding wounds does not depend on JNK signalling^15^. This suggests that parallel to JNK other signals are involved in syncytium formation after wounding.

In the adult fly, syncytium formation during would healing involves polyploidization and cell fusion, both dependent on Hippo/Yorkie signalling^16, 17^. While no connection with autophagy was shown in this system, it is worth noting that Yorkie can activate TORC1, a direct suppressor of autophagy, and that TORC1 is activated during larval wound healing, where it supports wound closure partly through S6K, one of its downstream effectors^6, 18^.

Whether autophagy and syncytium formation are causally connected is not known. However, during aging a progressive elevation of autophagy drives epidermal remodelling, morphological deterioration and formation of large multinuclear cells^19^. Global reduction of autophagy is associated with reduced plasma membrane degradation and aging of the epidermis^19^.

Here we directly tested the connection between autophagy and syncytium formation during wound healing and in the unwounded epidermis. We show that in *Drosophila* larvae elevated autophagy induces syncytium formation by membrane breakdown both in the epidermis and during epidermal wound healing.

## Results

### Autophagy and syncytium formation during wound healing

To analyse the function of autophagy for wound healing, we co-expressed a marker for autophagosomes (*GFP-Atg8a*) with RNA interference (RNAi) constructs against autophagy components in the epidermis and created epidermal wounds in early third larval instars. Atg8a is homologous to human MAP1 LC3 and localizes to all autophagic vesicles from the phagophore to the autolysosome^20^. In unwounded control epidermis, GFP-Atg8a is diffusely distributed in the cytoplasm and nucleus and is additionally seen in a few puncta in the cytosol (Fig. 1a). After wounding, new autophagosomes appear in the cells surrounding the wound^6^, increasing in number until completion of wound closure and dropping again to pre-wounding levels by 50-60 min after wound closure (Fig. 1a-c; Supplementary Video 1).

**Fig. 1.**
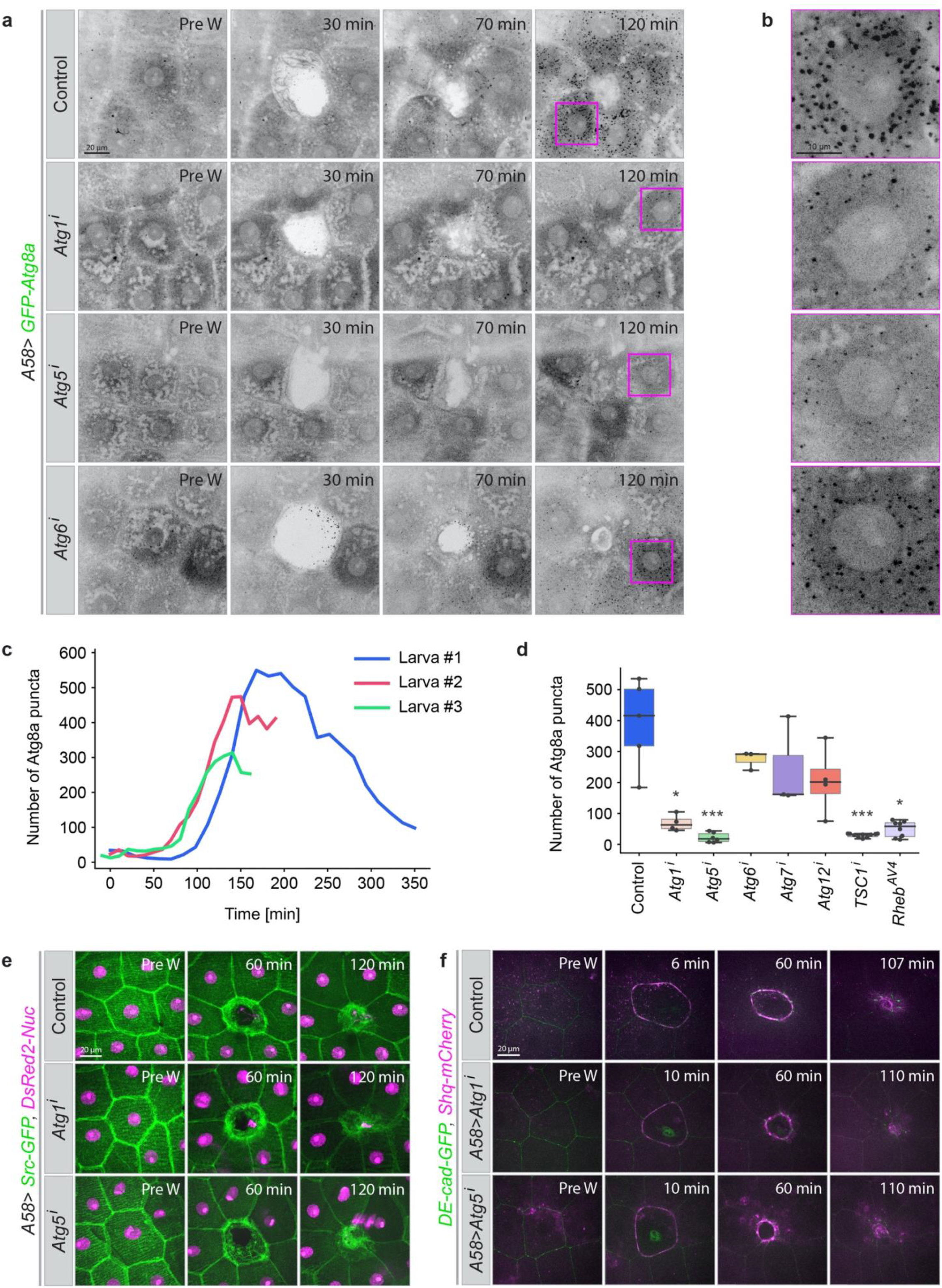
Autophagy during epidermal wound healing. **a-d,** Appearance of autophagic structures (marked with GFP-Atg8a) during wound closure in the epidermis of control third instar larvae and after epidermal knockdown of the autophagy pathway components Atg1, Atg5, Atg6, Atg7 or Atg12. All constructs are expressed in the epidermis under the control of the *A58-Gal4* driver. **a,** Time points from movies of wounded epidermis. The wounds have closed by 2 h in all cases. **b,** Higher magnification of the areas marked by magenta boxes at t=120 min. **c,** Quantification of the appearance of GFP-Atg8a puncta in the imaged area (10000 µm^2^) 3 control larvae, shown for each individual larva. **d,** Quantification of number of GFP-Atg8a puncta in different genetic conditions measured in an area of 10000 µm^2^ at the time of wound closure; n=4-10 larvae each genotype, for the detail see **Data analysis**. **e,f,** Effect of suppressing autophagy on wound healing and actin cable formation. Time-lapse series of single-cell wound healing in larvae expressing **e,** Src-GFP (green) and DsRed2-Nuc (magenta) to mark cell membrane and nuclei and **f,** endogenously GFP-tagged E-cadherin (DE-cad-GFP; green) and mCherry-marked myosin regulatory light chain (Sqh-mCherry; magenta) to visualize adherens junctions and actomyosin cables. **a,b,e,f,** z–projections of time-lapse series in early L3 larvae, n=9-15 larvae each genotype. Scale bars, **a,e,f,** 20 μm and **b,** 10 μm. Pre W: pre wounding. Images from **Supplementary Videos 1–3**. Genotypes of all images are listed in **Table 1**.

When autophagy was suppressed by knockdown of *Atg1* (the key component for autophagy initiation) or *Atg5* (a component for phagophore elongation) in the epidermis, the number of Atg8a vesicles in the cells around the wound was significantly reduced (Fig. 1a,b,d; Supplementary Video 1). Knockdown of *Atg6* (phagophore nucleation factor), *Atg7* or *Atg12* (phagophore elongation factors) also resulted in a decrease in autophagosomes, but the effect was much less pronounced, possibly because the RNAi constructs were less effective (Fig. 1a,b,d; Supplementary Video 1)

We analysed the effect of suppressing autophagy (using RNAi against *Atg1*, *Atg5, Atg6*, *Atg7* or *Atg12*) on the quality and rate of wound healing in larvae in which the plasma membranes were marked with Src-GFP or the adherens junctions with *DE-Cadherin-GFP* and the myosin regulatory light chain (MRLC) with *Sqh-mCherry* (a tagged form of *Drosophila* MRLC). Epidermal reduction of autophagy caused no morphological or developmental abnormalities in the epidermis and it did not affect the quality or rate of wound healing: lamellipodia, cell crawling, the assembly and contraction of the actomyosin cable, and the rate of wound closure were all normal and similar to controls (Fig. 1e,f; Supplementary Videos 2,3).

In the larva, the cells surrounding a wound form a large syncytium^13, 15^. To visualize syncytium formation in our experimental system, we created wounds in larvae with epidermal mosaics of cells expressing free cytosolic GFP (Fig. 2a,b). When we ablated an unmarked cell that was surrounded by both GFP-marked and unmarked cells, GFP gradually appeared in initially GFP-free cells around the time of wound closure (Fig. 2b; Supplementary Video 4). By the time the wound was fully closed, all cells surrounding the wound were GFP-positive, with an overall homogeneous GFP level. This behaviour was independent of the initial number of GFP-positive cells.

**Fig. 2.**
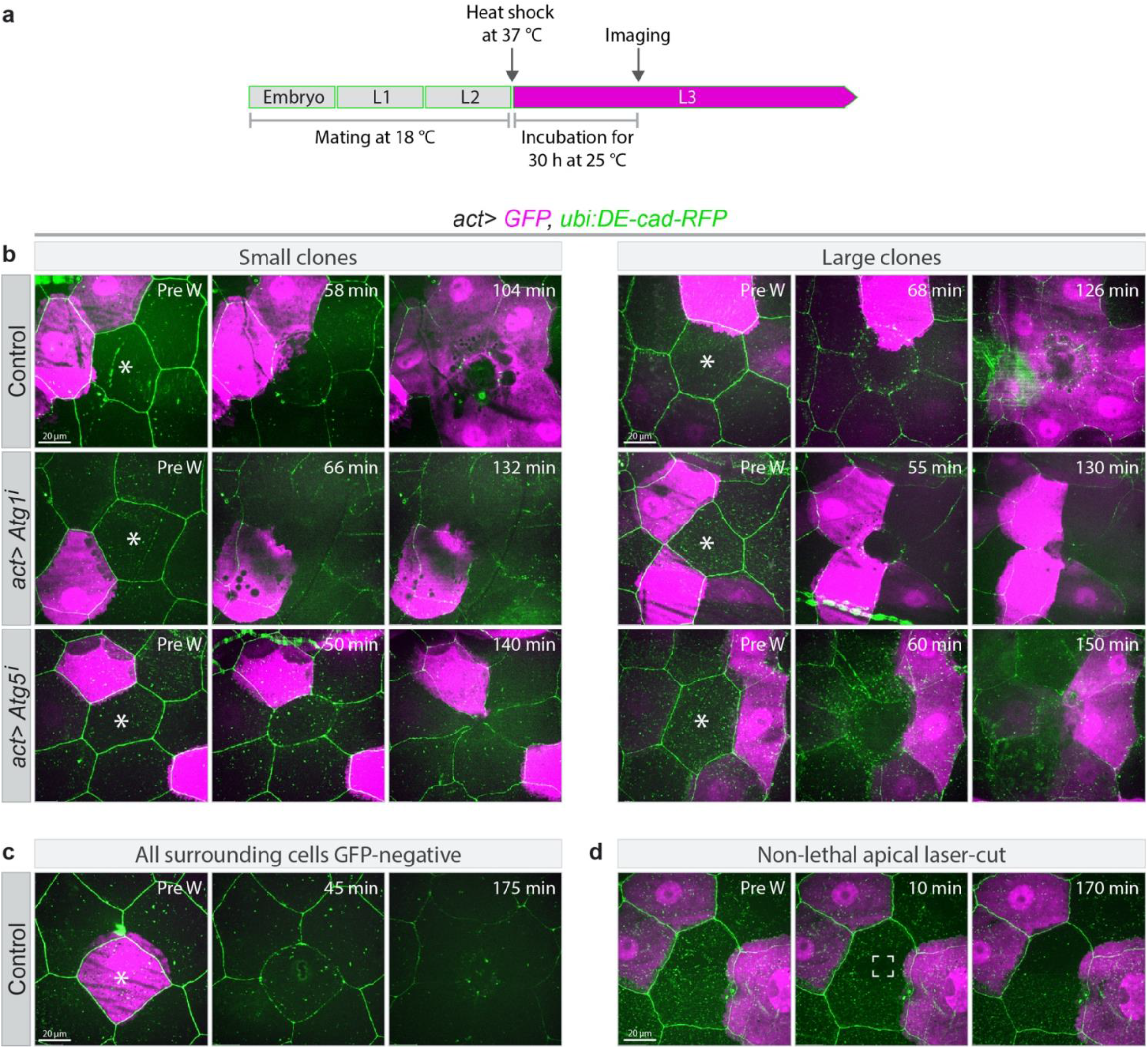
Syncytium formation during wound healing. **a,** Schematic for timing of transgene expression and start of live imaging. Gene expression was induced at the end of the second larval instar, laser ablation and imaging started 30h later. **b,** Wound healing in epithelia with clonally expressed cytoplasmic GFP (magenta) under the *actin5c-Gal4* driver in control larvae (*act>GFP*) or larvae expressing RNAi constructs specific for *Atg1* or *Atg5*. Laser ablated cells are marked with white asterisks. To visualize cell borders DE-cad-RFP (green) was expressed in all tested genotypes. By the end of wound closure, GFP from the clonal cells has spread to all cells around the wound in the control, but not if autophagy is suppressed, regardless of the number of cells initially expressing GFP. **c,** Control larva in which a GFP-expressing cell was wounded. No GFP is induced in or taken up by the surrounding cells. **d,** Control experiment in which the central cell was damaged but not killed (white marked area). No wound response occurs and no GFP-leakage between neighbouring cells is seen. n=6–9 larvae each genotype. The control pre-wounding small clone in b (top left panel) is from Kakanj et al. 2016. **b,c,d,** z–projections of time-lapse series. **b-d,** Scale bars, 20μm. Pre W: pre-wounding. Images from **Supplementary Videos 4–7**.

We confirmed that the appearance of GFP in the initially GFP-negative cells was not due to *de novo* synthesis in response to wounding. When we ablated a cell that was surrounded by cells without GFP we observed no GFP expression during an imaging time of ∼3 hours (Fig. 2c; Supplementary Video 5a). When we only damaged the apical membrane of a cell without fatally wounding it, the GFP-negative cells surrounding the damaged cell remained GFP-negative (Fig. 2d; Supplementary Video 5b). This indicates that our experimental manipulations do not induce *de novo* GFP expression in GFP-negative cells around the wound. Instead, we conclude that the GFP is redistributed from GFP-expressing to GFP-negative cells, and that the wound healing response includes a process by which the cells around the wound gradually share their cytoplasm.

One possible explanation for the observations so far is that the redistribution of cytoplasm and the induction of autophagy observed during wound healing are causally related. We tested this by ablating cells in epithelia in which *Atg1* or *Atg5* were knocked down and GFP expressed in single or multiple clonal cells. We saw no intercellular redistribution of GFP in these experiments (Fig. 2b; Supplementary Videos 6,7). This shows that cytoplasmic mixing and formation of a syncytium depend on autophagy and that autophagy is required in a cell-autonomous manner for a cell to become part of the syncytium.

### Effect of Autophagy on epidermal morphology

We also analysed the effect of autophagy induction in the unwounded epidermis. The basal level of autophagy is low and only few Atg8a-positive vesicles are detected in the epidermis of control larvae (Fig. 3a,b). We activated autophagy artificially with two different overexpression constructs for Atg1, Atg1S (‘strong’) and Atg1W (‘weak’) (**Table 2** and Scott et al. 2007). Both led to an increase in the number of Atg8a vesicles (Fig. 3a-c). We also observed an increase of vesicles marked with LAMP1-GFP, an endo-lysosomal marker, indicating that the entire autophagic pathway was active (Supplementary Fig. 1a,b).

**Fig. 3.**
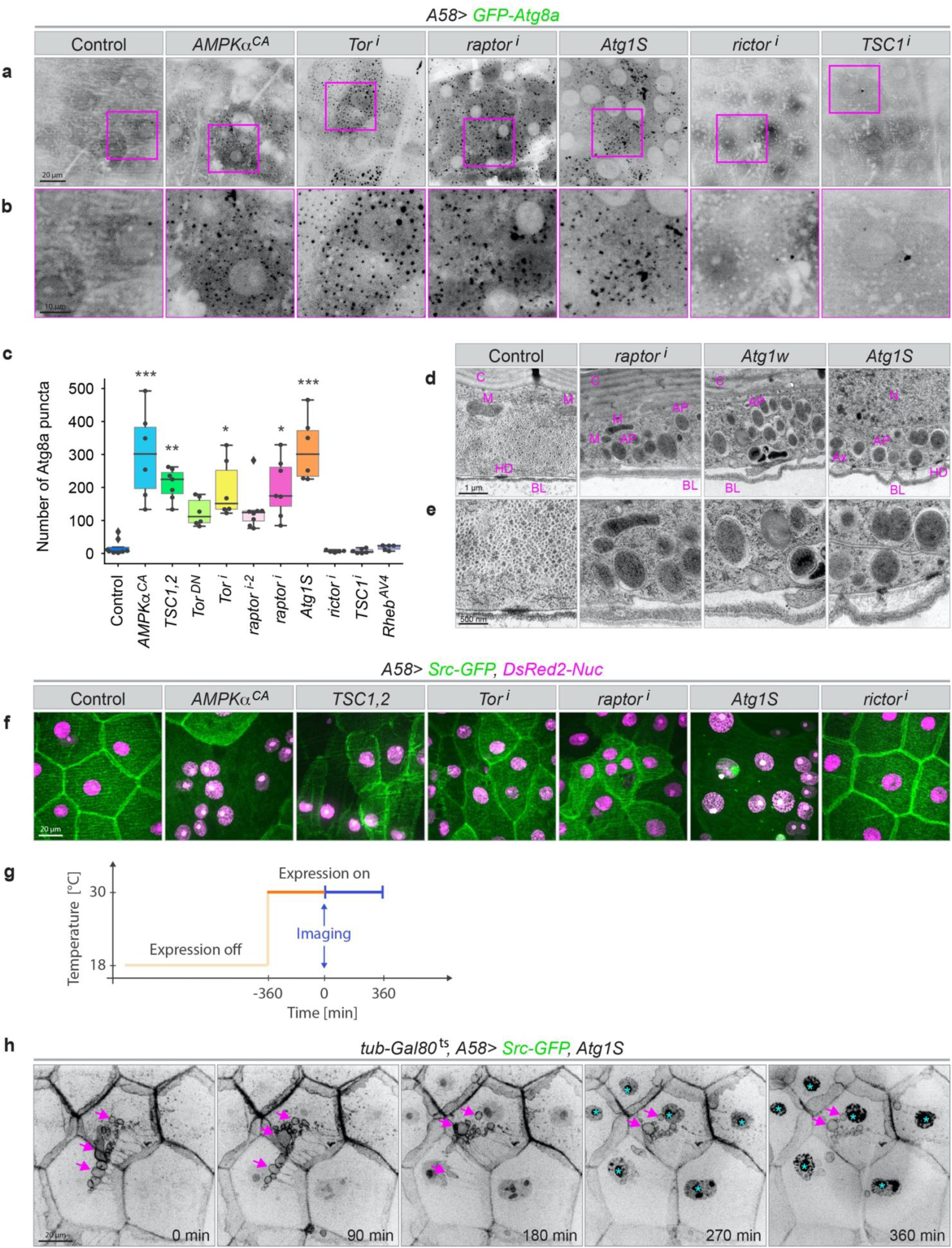
Autophagy in unwounded epidermis. **a-d,** Control of epidermal autophagy by TOR signalling. **a,** Epidermis of third instar larvae expressing the autophagosome marker GFP-Atg8a together with constructs for up-or down-regulating the autophagy pathway in the epidermis. Healthy epidermis contains few autophagosomes, but artificially activating autophagy through overexpression of Atg1 or blocking TOR signalling leads to accumulation of autophagosomes. **b,** Higher magnification of the areas marked by magenta boxes in (**a**). **c,** Quantification of Atg8a puncta in an area of 10000 µm^2^, n=6–8 larvae each genotype. **d,e,** Transmission electron micrographs of sections through the epidermis of larvae with elevated autophagy at two different magnifications. AP: autophagosome; Ax: cross-section of an axon; BL: basal lamina; C: cuticle; HD: hemidesmosome; M: mitochondrion; N: nucleus. **f,** Disruption of epidermal morphology after upregulation of autophagy. The membrane marker Src-GFP is lost from large areas of the epidermis and nuclei have lost their regular spacing. **g,** Schematic for temporally controlled transgene expression and imaging in (**h**). Gene expression is induced at the end of the second larval instar, live imaging started 6 h later and continued for an additional 6 hours. **h,** Example of membrane dynamics after time-controlled Atg1S expression. Src-GFP containing material appears to be taken out of and eventually detached from lateral cell membranes (arrows). Over the period of observation, abnormal accumulation of GFP is seen in the nuclei (cyan asterisks). t = 0 is 6h after A58-Gal4 activation. Image from **Supplementary Video 8**. **a,b,f,h,** z– projections; n=20-50 larvae each genotype. Scale bars: **a,f,h,** 20 μm, **b,** 10 μm, **d,** 1 μm and **e,** 500 nm.

Autophagy is normally kept inactive via TORC1 signalling, and we tested the effect of both disruption and activation of TORC1 on autophagy. Experimental activation of TORC1, either through overexpression of an upstream activator, *Rheb^AV4^*, or knockdown of the TORC1 inhibitor, *TSC1* (*TSC1^i^*), had little effect on the number of Atg8a-positive vesicles in the tissue (Fig. 3a-c). In contrast, expression of the TORC1 antagonists, *AMPKα^CA^*, *TSC1*, *TSC2*, or a dominant-negative version of Tor (*Tor^DN^*), or downregulation of *Tor* or *raptor* by RNAi increased the number of Atg8a-positive and LAMP1-positive vesicles (Fig. 3a-c; Supplementary Fig. 1a,b). Epidermal knockdown of *rictor*, a component of TORC2, did not induce autophagy (Fig. 3a-c; Supplementary Fig. 1a,b). Thus, the suppression of autophagy in the larval epidermis depends specifically on TORC1.

Transmission electron microscopy (TEM) images showed large numbers of autophagosomes and autolysosomes after TORC1 reduction or Atg1 overexpression while very few were present in control epidermis confirming that the increase in Atg8a- and LAMP1 puncta corresponded to an increase in autophagosomes and autolysosomes (Fig. 3d,e).

The increase in Atg8a-positive vesicles caused by overexpression of Atg1 was always similar in quality but stronger in extent than that resulting from reducing TORC1 activity directly or through its upstream regulators.

In the experiments above we noticed that cells often appeared morphologically abnormal. We then used the plasma membrane marker Src-GFP to analyse this effect. Reduction of TORC1 activity or expression of Atg1 led to distortions of epidermal morphology, with effects on the shape and orientation of cells, localisation of the nuclei, and size heterogeneity in cells and nuclei (Fig. 3f). Most strikingly, cell outlines often appeared to be interrupted or missing altogether (Fig. 3f). Time-controlled transgene activation using the temperature-sensitive Gal80^ts^ system showed that defects were visible within 6 hours of TORC1 depletion or activation of Atg1 (the earliest possible time point for analysis, since the marker constructs were not detectable sooner). These defects became exacerbated over the following 18 hours, with a progressive change in organisation of lateral membranes, accompanied by the formation of large vesicles that moved toward the centre of the cell and then disappeared (Fig. 3g,h, Supplementary Fig. 1c; Supplementary Video 8). Together, these results indicate that disruption of TORC1 signalling causes exacerbated autophagy and altered epidermal morphology.

### Autophagy and the formation of epithelial syncytia

The results so far could mean that the missing cell outlines were due to a change in membrane properties, so that Src-GFP could no longer be recruited, or there could be gaps or defects within the membranes. To distinguish between these possibilities, we first examined other membrane markers. The main components of the apical cell polarity complex Par3 (Bazooka) and aPKC, normally located in the apical domain of the lateral plasma membrane, and the septate junction proteins Fasciclin III (FasIII) and Neuroglian (Nrg) all showed abnormal distributions or were absent over large areas in TORC1-depleted and Atg1-expressing epithelia (Fig. 4a-c; Supplementary Fig. 2a-d). Both live imaging with GFP-constructs and immunofluorescence of fixed material gave the same results, showing that mis-localisation of GFP or absence of determinants was neither due to breakdown of GFP nor to fixation problems.

**Fig. 4.**
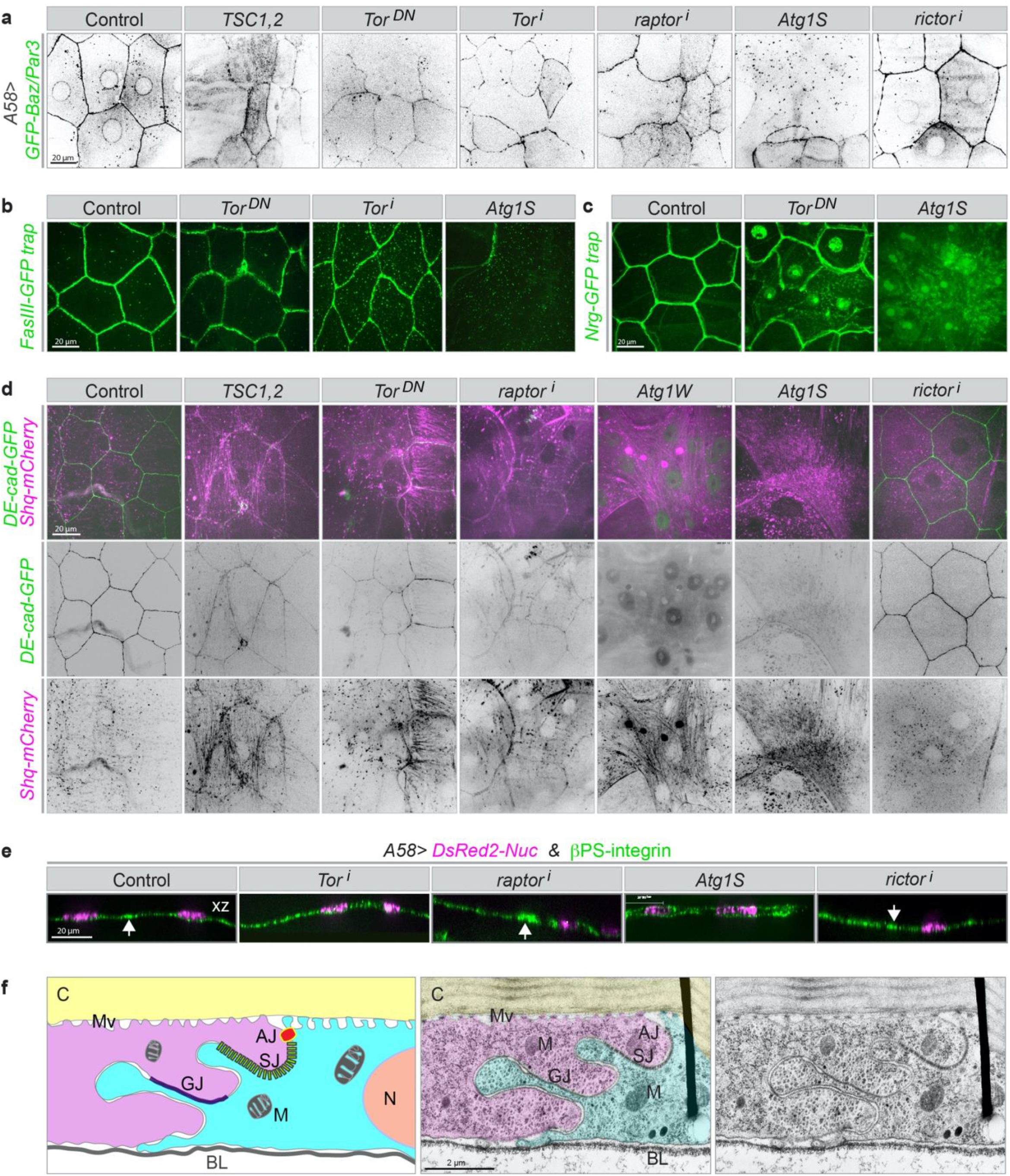
Effect of uncontrolled autophagy on plasma-membrane associated proteins. **a-e,** Surface views (**a-d**) and z-sections (**e**) of third instar larval epidermis expressing the indicated fluorescent markers and RNAi or overexpression constructs. **a,** Bazooka/Par3, normally seen at the apical adherens junction and in a perinuclear position, is lost or reduced in large areas under uncontrolled autophagy as are lateral membrane markers, **b,** Fasciclin-III (FasIII), **c,** neuroglian (Nrg) and **d,** adherens junctions (DE-cad), whereas large bundles decorated with MRLC appear. **e,** The baso-lateral transmembrane protein β-integrin is not lost; in extreme conditions, it is seen in both the apical and basal membranes (overexpression of Atg1S). Arrows in the z-sections point to high accumulation of integrin along the folded lateral membranes. **f,** Electron micrograph of a section through the larval epidermis to show the highly folded lateral junction between two epidermal cells (magenta and cyan). Left, cartoon; middle, false colouring; right, original image. C: cuticle; M: mitochondrion; BL: basal lamina; AJ: adherens junction; SJ: septate junction; GJ: gap junction, N: nucleus, Mv: apical microvilli connecting the cuticle to the cell. **a-e** n=15–40 larvae each genotype. Scale bars: **a-e,** 20 μm and **f,** 2 μm. Images from **Supplementary Videos 9–11**.

To assess the extent of cell abnormality we also imaged other molecules involved in cell shape and polarity. We found that adherens junctions (visualized with DE-Cadherin-GFP) were reduced or absent, while myosin (MyoII, visualized with the mCherry tagged non muscle type II myosin regulatory light chain, Sqh-mCherry) had formed strong cortical and cytosolic filaments (Fig. 4d). Experimental activation of TORC1 had the opposite effect: it reduced myosin level similar to knockdown of RhoA or Rok (Supplementary Fig. 2e).

TORC2 is involved in regulation of plasma membrane and actin cytoskeleton homeostasis both in yeast and mammalian cells^21, 22^, but reduction of TORC2 activity had no effect in any of the assays we used for autophagy or cell morphology (Fig. 3a-c,f, 4a,d; Supplementary Fig. 1a,b).

Z-sections of Src-GFP expressing cells showed that the basal and apical membranes appeared intact (Supplementary Fig. 3a). We stained also for the basal marker β-integrin, which is normally localised at the basal and basolateral plasma membrane up to the border of the septate junctions (Fig. 4e; Supplementary Fig. 3b). Because the lateral membrane is highly folded (Fig. 4f), the outlines of cells are seen as a fuzzy band of increased integrin staining. The epithelium is very thin and the nuclei extremely flattened, but in spite of the low resolution, z-sections show the integrin signal in a thin line below the nuclei in control epidermis. This separation is less clear in the absence of TORC1 signalling, and strongly disturbed when Atg1 is overexpressed. In this case, integrin is seen both below and above the nuclei, suggesting that it is also present in the apical compartment (Fig. 4e).

The basal membrane is not only present but it is also intact, as shown by the fact that we see no significant loss of cytosolic GFP from the epidermis, either in untreated or in wounded epithelia (see below). This may also explain why these larvae can survive throughout larval development.

Finally, despite defects in the lateral plasma membranes, the abnormal epithelia resulting from uncontrolled autophagy were able to form an actomyosin cable around wounds. While the wounds healed more slowly, they eventually closed completely, and most of the larvae (∼75%) survived to the pupal stage (Supplementary Fig. 4a,b; Supplementary Videos 9-11). Thus, the defects in morphology are not simply a sign of the epithelium disintegrating or dying, but the epithelium is viable and physiologically active.

The results so far indicate that activation of autophagy results in the disappearance of lateral membrane markers. If this is due to the loss of lateral membrane then cytosolic proteins should be able to diffuse between cells. We tested this with fluorescence loss in photobleaching (FLIP) to monitor the intercellular movement of GFP. In control experiments we bleached the GFP in an area of 179 µm^2^ within a cell for 3 min, resulting in removal of GFP not only from the bleached area but also from the entire cell (Fig. 5a-c; Supplementary Video 12). The GFP in neighbouring cells was not affected, and no GFP re-appeared in the bleached cell over the next 20 min. In contrast, a 3-min illumination of an area of 179 µm^2^ in larvae expressing Atg1 in the epidermis was insufficient to remove GFP from the marked area (Fig. 5a-c; Supplementary Video 12). Photobleaching of a larger area (1098 µm^2^; 1-4 nuclei) led to removal of GFP from this area, but the area remained dark only for a few seconds, after which the fluorescence signal re-equilibrated to the same level as the surrounding area, which we interpret as diffusion of GFP into the bleached region from neighbouring areas (Fig. 5a-c; Supplementary Video 12). Z-sections of movies showed clear diffusion of GFP laterally within the epithelium, and never any basal or apical leakage (Supplementary Video 13).

**Fig. 5.**
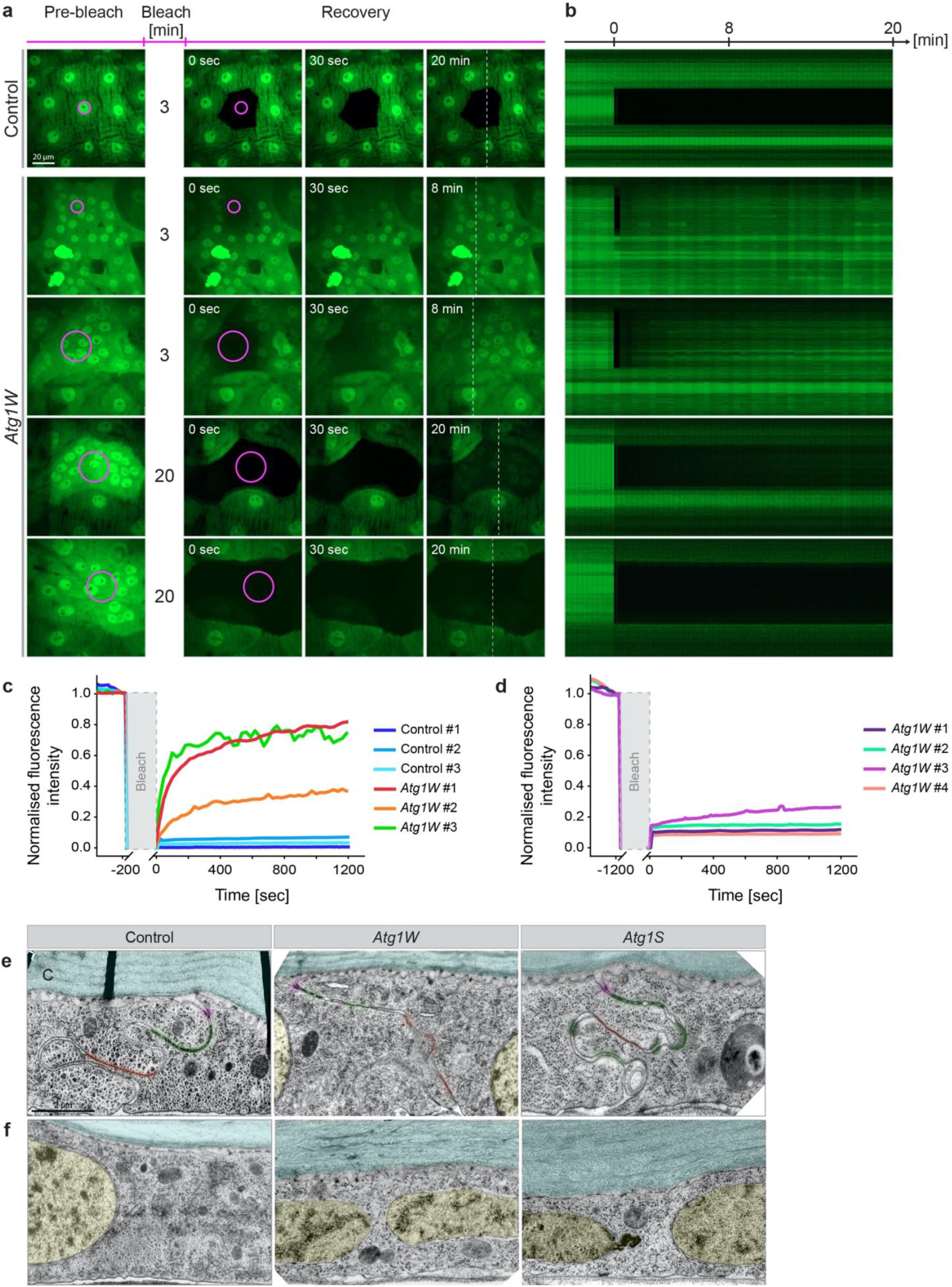
Loss of lateral membrane integrity. **a,b,** Fluorescence loss in photobleaching (FLIP) to test free cytoplasmic GFP motility within the epidermis. A small area (magenta circle; 179 or 1098 µm^2^) was laser-illuminated for the indicated times (3 or 20 min) in control or Atg1-expressing epidermis also expressing free GFP. **a,** Snapshots before bleaching and at 3 points of recovery. **b,** Kymographs along the broken line in (**a**) during recovery. **c,d,** Quantification of fluorescence recovery after bleaching, shown separately for 3 and 20 min bleaching protocols**. e,f,** Electron micrographs showing morphological defects or absence of lateral cell membranes in epithelia with upregulated autophagy. **e,** Membrane domains with tight apposition between neighbouring cells are marked in magenta (adherens junctions), green (septate junction) and orange (gap junctions). **f,** Nuclei (yellow) and lateral membranes in healthy epidermal cells cannot be shown in one image because they are too far apart, whereas when autophagy is upregulated, nuclei are often found close together and not separated by plasma membranes. C: cuticle (blue). **a-d** n=5-9 larvae for each FLIP protocol. Scale bars: **a,** 20 μm and **e,f,** 2 μm. Images from **Supplementary Videos 12–14.**

To assess the extent of the region connected to the bleached area we increased the time of photobleaching to 20 min (Fig. 5a,b,d; Supplementary Video 14). This led to an ∼85% reduction of the GFP signal in a much larger area (7000-15000 µm^2^, with 10-24 Nuclei), showing that the epidermal cells in this patch had formed an extended syncytium from which GFP entered the area being bleached. In most cases the fluorescence signal did not recover over the next 20 min, showing the syncytium had been exhausted of GFP and was separated from other parts of the epidermis. Thus, activation of autophagy induces the formation of syncytial patches in the unwounded epidermis.

Furthermore, transmission electron micrographs showed that lateral membranes were missing in many places (as judged by adjacent nuclei not being separated by a membrane), but the basal membrane was intact (Fig. 5e,f). The epithelium also showed a range of other defects, from an abnormal basal lamina and apical cuticle to deformed nuclei (Supplementary Figs. 5a,b, 6b,c).

The effect of uncontrolled autophagy on cell morphology was not unique to the epidermis. The ectodermal driver also directs expression in the tracheal system, and the tracheal trunk and branches were severely distorted. Similarly, the enterocytes of the gut and secretory cells of salivary glands lost lateral membrane markers and had abnormal shapes. Yet, as in the epidermis, the epithelial barrier function was not compromised.

Together, these experiments indicate that activation of autophagy affects the lateral membrane of the epidermal cells leading to the formation of syncytia in the unwounded epidermis, which appears reminiscent of the syncytium observed during wound healing. Both loss of TORC1 or hyperactivity of Atg1 disrupt cell integrity in a specific manner: lateral membranes are lost, but basal and apical membranes remain intact but adopt inappropriate identities. As a consequence, autophagy must be kept repressed in epithelia for cell integrity and polarity to be retained, and this repression requires TORC1 activity.

### TORC1-dependent autophagy alleviates nuclear defects in models of laminopathy

To investigate whether other membranes were affected by uncontrolled autophagy we used fluorescent markers for the mitochondria (mito-GFP), ER (KDEL-RFP), Golgi (GalT-GFP) and nuclear envelope (endogenously GFP-tagged Kugelkern (Kuk), a *Drosophila* nuclear lamina protein). Reduction of TORC1 or expression of Atg1 had no apparent effect on the number or distribution of the mitochondria (Supplementary Fig. 5c). This fits with TEM images, which show a normal density and morphology of mitochondria. However, the number of KDEL- and GalT-positive puncta was reduced, suggesting that they were also being depleted by autophagy (Supplementary Fig. 5d-g).

Nuclei marked with GFP-Kuk in control cells appear round with a homogeneous GFP signal at the lamina (Fig. 6a). Loss of TORC1 or Atg1 hyperactivation caused abnormalities in nuclear morphology and shape, disruption of the nuclear membrane with irregular GFP distribution and vesicular structures inside and outside of the nucleus (Fig. 6a; Supplementary Fig. 6a). Other markers associated with the nuclear lamina were also abnormal. Bazooka/Par3, normally localised at the nuclear envelope, was not detectable when autophagy was activated in epidermal cells (Fig. 4a; Supplementary Fig. 2a). TEM imaging supported these observations, in addition showing autophagosomes within the nucleus (Supplementary Fig. 6b,c). The defects were not restricted to nuclear morphology but extended to chromatin organisation: an RFP-tagged version of His2Av (*endo:His2Av-mRFP1*) revealed abnormal chromatin condensation in cells with upregulated autophagy (Supplementary Fig. 6d; Supplementary Video 15).

**Fig. 6.**
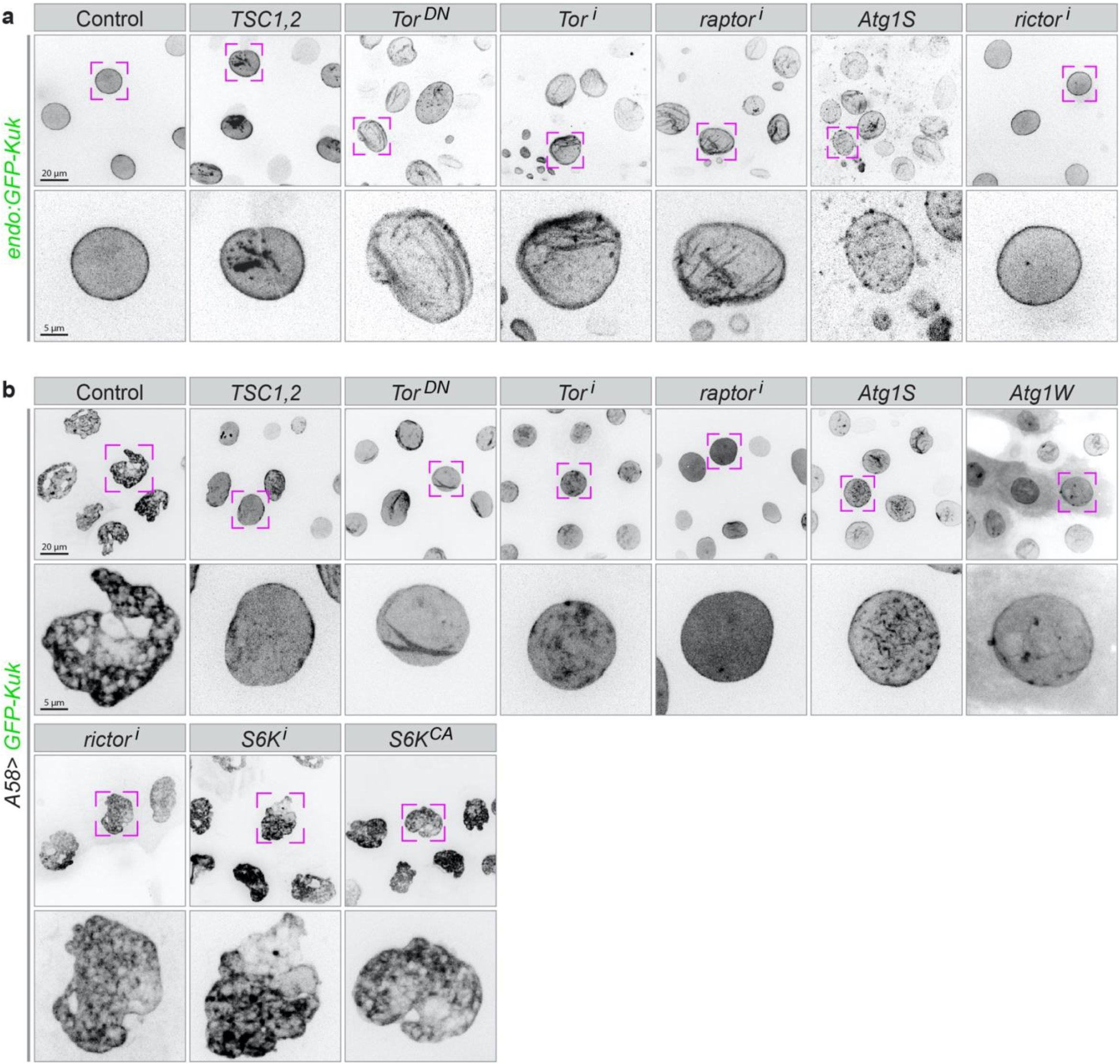
Effect of uncontrolled autophagy on nuclear morphology. **a,** The nuclear lamina is visualized using a *GFP-tag* inserted into the endogenous locus of the *Kugelkern (Kuk)* gene in animals expressing the indicated overexpression or RNAi constructs in the epidermis. **b,** A GFP-tagged transgenic construct of *Kugelkern* is co-expressed with the indicated overexpression or RNAi constructs. High levels of Kuk induce lobulation and other nuclear defects, which are ameliorated if TOR is downregulated or autophagy upregulated, but not if other branches of the TOR signalling pathways (S6K or rictor) are modified. The lower row shows higher magnification of the nuclei marked above. n=20–30 larvae each genotype. Scale bars: **a,b** upper rows, 20 μm; lower rows, 5 μm.

Defects in the nuclear lamina are associated with a spectrum of diseases known as laminopathies^23–26^. In a *Drosophila* model for nuclear laminopathy overexpression of Kuk or Lamin B (LamB) leads to the lobulation and increased folding of the nuclear envelope in the muscle and a reduced lifespan^25, 27–29^. Overexpression of GFP-Kuk or RFP-LamB in the epidermis caused also an abnormal morphology of nuclei but the larvae developed to fertile adult flies, without visible defects (Fig. 6b; Supplementary Fig. 6e). Induction of autophagy by TORC1-reduction or Atg1 overexpression largely reduced the nuclear defects caused by elevated levels of Kuk or LamB (Fig. 6b; Supplementary Fig. 6e), pointing to the possibility of the autophagy machinery competing for the membrane needed to increase the nuclear envelope. Conversely, this shows that elevated autophagy likely also depletes the nuclear envelope, possible via the ER. Thus, autophagy can counteract the nuclear defects seen in this laminopathy model.

### Relevance of autophagy for plasma membrane integrity

We have seen that reduction of TORC1 induces autophagy and causes similar defects in epidermal cells as the more direct activation of autophagy by Atg1 does. Next, we tested whether TORC1 acts exclusively through inducing autophagy, and if so, whether the entire autophagic process has to be completed for the membrane defects to occur. Apart from repressing Atg1 activity, TORC1 also activates S6K. Constitutive activation of S6K (*S6K^CA^*) did not improve any aspect of cellular defects caused by the TORC1 block, indicating that loss of S6K activity is not responsible for the defects caused by loss of TORC1 (Supplementary Fig. 7a). TORC2 reduction also did not improve any of the cellular defects derived by TORC1 reduction (Supplementary Fig. 7a).

To assay the different steps of the autophagic pathway we first tested the efficiency of RNAi constructs in suppressing autophagy in the epidermis. As during wound healing, knockdown of *Atg1* or *Atg5* both reduced the number of Atg8a-positive vesicles in TORC1-depleted epidermis to control levels (Supplementary Fig. 7b,c). In contrast, knockdown of *Atg6* or *Atg7* reduced the number of Atg8a-positive vesicles only slightly, similar to their limited effect on autophagosomes during wound healing (Supplementary Fig. 7b,c and Fig.1d). These two were therefore not suitable as tools to block later stages of autophagy in this system.

To test whether the effects of TORC1 on cell membrane were caused exclusively through the release of Atg1 activity and the autophagy pathway, we down-regulated *Atg1* or *Atg5* together with TOR or raptor. In these larvae, the membrane defects caused by uncontrolled autophagy were largely abolished, except in a few cells at segment borders (Fig. 7a; Supplementary Fig. 8a). The suppression of the effects of TORC1 knockdown is not due to titration of Gal4 activity because other constructs such as *UAS-rictor^i^* did not have this effect (Supplementary Fig. 7a). This shows that Atg1 is needed to mediate the effects of TORC1 on lateral membranes. Similarly, downregulation of *Atg5* suppressed the effects of both TORC1 reduction and Atg1 hyperactivity (Fig. 7a; Supplementary Fig. 8a). We also confirmed with the FLIP assay that syncytium formation was reduced in Atg1W, Atg5i larvae. GFP was no longer able to move into bleached areas from large surrounding regions; instead, only one cell was bleached, as in controls, or at most one neighbouring cell also lost fluorescence (Fig. 7b-d; Supplementary Video 16). Blocking autophagy also re-established the proper distribution of DE-cadherin, integrin and MyoII (Fig. 8a,b; Supplementary Fig. 8b).

**Fig. 7.**
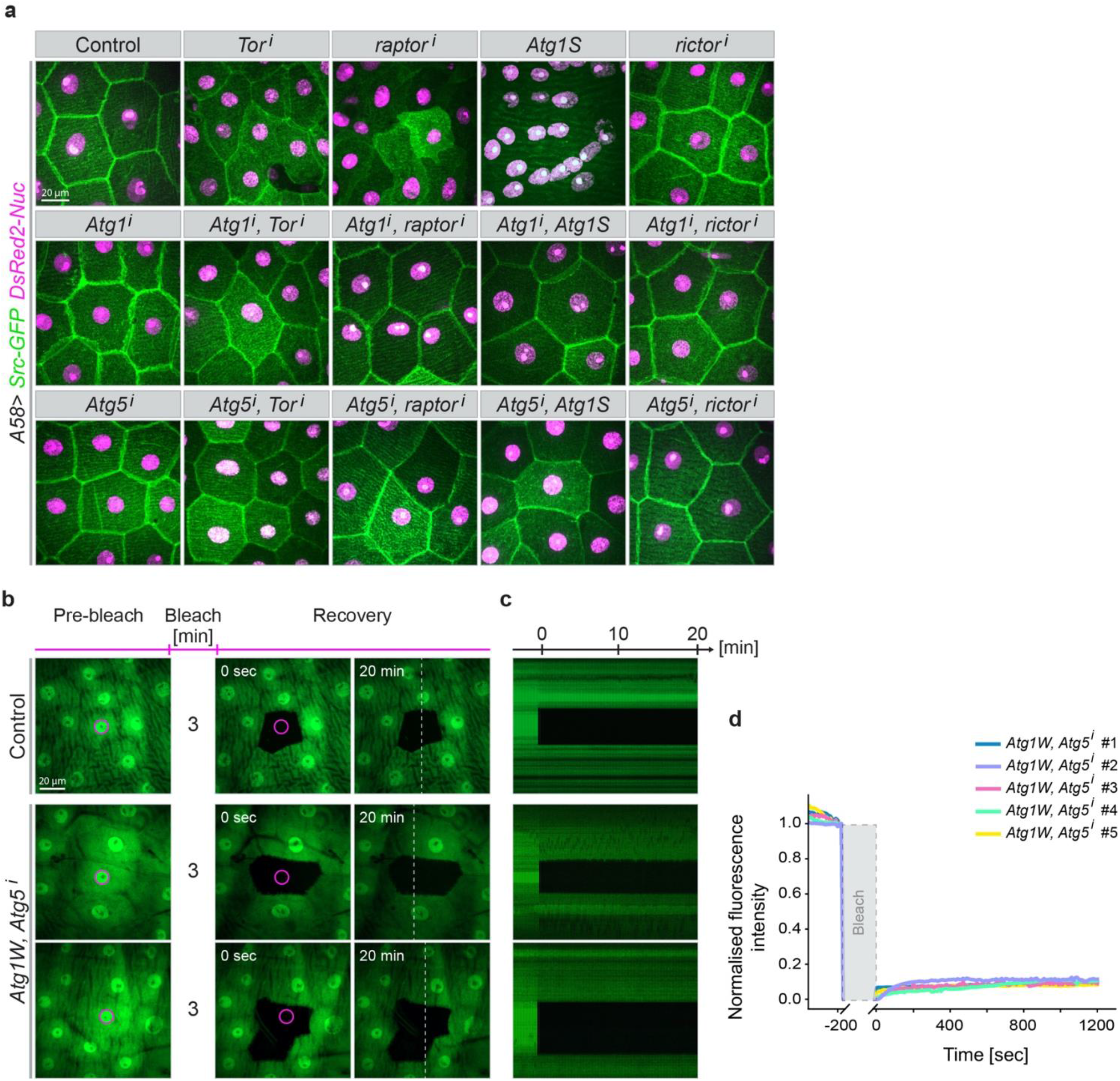
Epistasis of TOR and autophagy. **a,** The indicated overexpression and RNAi constructs were co-expressed with markers for the plasma membrane (Src-GFP, green) and nuclei (DsRed2-Nuc, magenta) in the larval epidermis. The effects of upregulating autophagy (top row) are suppressed when autophagy is blocked by simultaneously downregulating Atg1 or Atg5. n=20-40 larvae each genotype. **b-d,** Atg1-induced syncytium formation is abolished when Atg5 function is downregulated (compare to rows 2 and 3 in Fig. 5 **a,b)**; same representation as shown in Fig. 5a-d. Scale bars: **a,b,** 20 μm. Images from **Supplementary Video 16.**

**Fig. 8.**
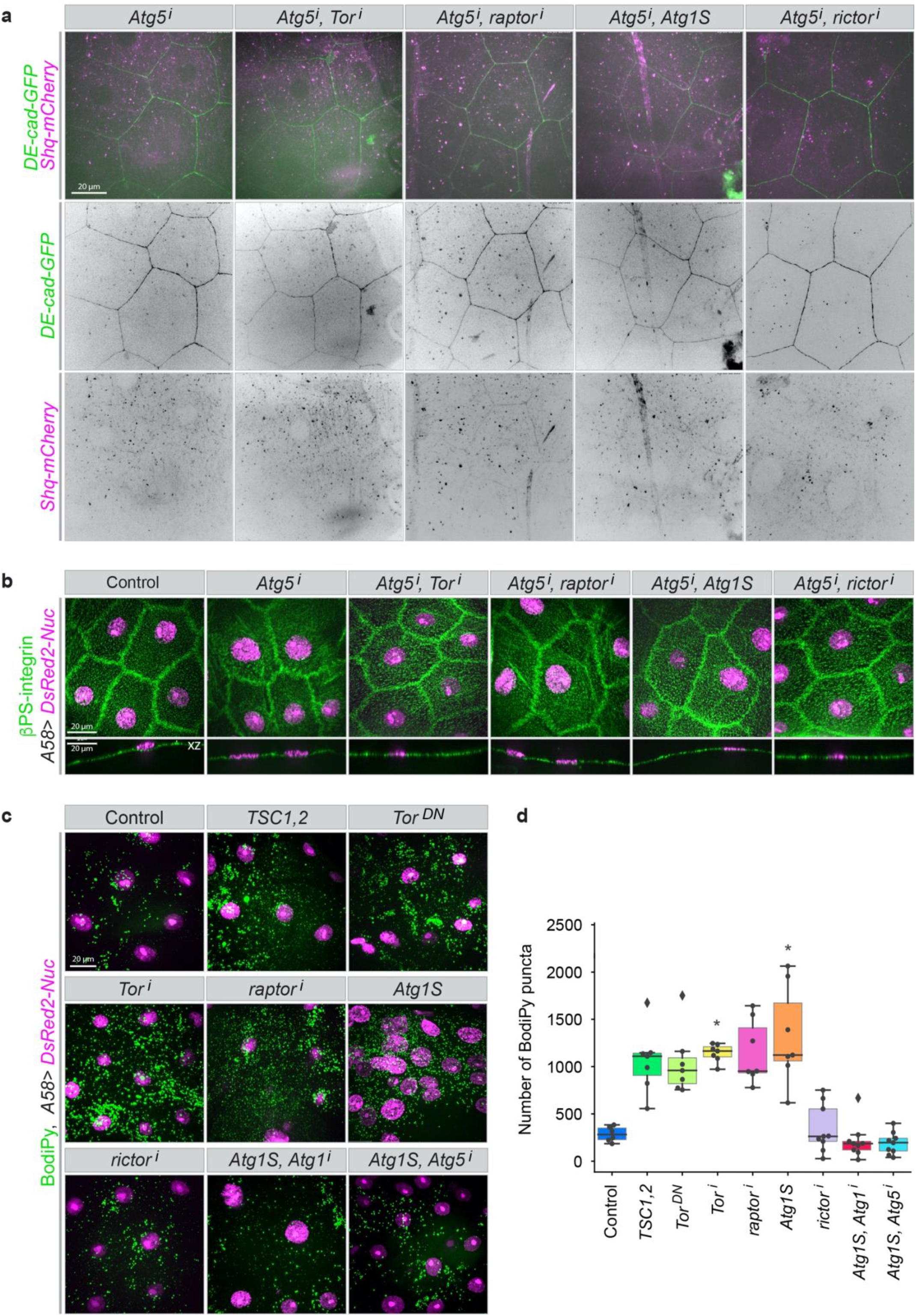
Dependence of cellular defects on functional autophagy. **a,b,** Blocking autophagy by downregulating Atg5 suppresses the cellular defects caused by down-regulation of TOR signalling: **a,** loss of lateral membranes, formation of actin bundles; **b,** integrin mislocalisation (compare to **Fig. 4 d,e and Suppl. Fig. 2e and 3b**). **c,d,** Accumulation of neutral lipid in the cytoplasm under conditions of uncontrolled autophagy. **c,** Fixed epidermis from larvae expressing the indicated constructs was stained with BodiPy 493/503 to mark lipid accumulations. **d,** Quantification of the number of lipid puncta in an epidermal area of 10000 µm^2^ in the indicated conditions. **a-d,** n=7–20 larvae each genotype. Scale bars: 20 μm. Images from **Supplementary Video 18.**

In contrast, a chloroquine-block of the last step of autophagy, i.e. the fusion of the autophagosomes with lysosomes, resulted in the expected accumulation of Atg8a-puncta but did not suppress the removal of lateral membrane resulting from TORC1 depletion or Atg1 activation (Supplementary Fig. 9a-c; compare to **Fig. 3 a-c**). This shows that the removal of lateral membrane requires the formation of autophagosomes but not the completion of autophagy, i.e. the final degradation of material engulfed by autophagosomes. It also suggests that the lateral plasma membrane material is incorporated into autophagosomes. Staining with the lipophilic dye BODIPY 493/503 confirmed the strong accumulation of lipids in cytoplasmic vesicles in TORC1-depleted epithelia and other conditions where the lateral membranes were disrupted (Fig. 8c,d).

Activation of autophagy also affected the distribution of actomyosin. We tested whether this was an independent, parallel effect, or part of the mechanism to remodel the plasma membrane. To examine this, we interfered with MyoII activation with a dominant negative version of the MyoII activator Sqa^T279A^ (“Sqa^KA^“) or RNAi against *Rok*. These treatments led to the disappearance of the abnormal actomyosin fibres^30^, but did not suppress the membrane defects (Supplementary Fig. 9d,e, 2e). Thus, abnormal MyoII activation is not responsible for the autophagy-induced membrane defects.

The fact that elevated Atg1 activity was sufficient to induce the abnormalities suggests that they do not depend on other TORC1-dependent signalling events. We directly tested whether these defects depended on autophagy by reducing Atg1 or Atg5 levels in the background of TORC1 reduction or Atg1 over-activation. This significantly restored wound healing (Supplementary Fig. 10a,b; Supplementary Videos 17,18). The actin cables took 16–30 min instead of 40–60 min to form (normal epidermis: 10 –12 min) and they were less thick. The total time to complete wound closure was similar as in normal epithelia (120 ± 20 min, depends on wound size). We had seen above that knockdown of Atg1 is not sufficient to completely suppress the effects of Atg1 overexpression on epithelial cell integrity. Therefore, here too, the remaining defects can probably be ascribed to a remaining slightly elevated level of Atg1 activity, both after TORC1 knockdown and Atg1 upregulation. Atg1 has been observed to affect phosphorylation of the activator, Sqa, and could thereby potentially directly modulate actomyosin in our experiments^30^. However, the finding that knockdown of Atg5 is able to suppress the effects on the actomyosin and wound healing shows that the defects do not result from a direct action of Atg1 on myosin but again require the autophagic pathway to occur. They therefore seem to be an indirect effect of membrane being sequestered in autophagosomes.

## Discussion

Our findings identify: i) a novel role for the TORC1/autophagy pathway in controlling plasma membrane integrity and homeostasis, ii) autophagy as a necessary and sufficient inducer of syncytium formation in the epithelium and during wound healing, iii) wounding as a trigger for autophagy and iv) the plasma membrane as potential source for autophagosome formation. On the basis of our observations we discuss the role of autophagy for syncytium formation, epithelial barrier function, phagocytosis, actomyosin organisation and homeostasis of endomembranes.

Multinucleated cells form during development and aging or under stress conditions in diverse tissues and organisms by different mechanisms such as cytokinesis failure, entosis, cell-cell fusion or by lateral membrane breaching^31, 32^.We will concentrate only on the latter here, and in particular on the connection to autophagy.

A number of studies indicate a correlation between autophagy and syncytium formation. The best studied cases are the fusion of myoblasts during muscle fiber formation and of trophoblast cells during placenta formation^33, 34^. In adult mouse the stem cells of skeletal muscles (myosatellite cells) are quiescent until external stimuli such as exercise or injury trigger their re-entry into the cell cycle and their differentiation to muscle progenitor cells (myoblasts). Myoblasts which have differentiated from adult muscle stem cells after external stimuli such as exercise or injury differentiate and fuse to form new multinucleated myofibers. TORC1-dependent autophagy is induced during myogenesis in physiological and pathological conditions such as, fasting, atrophy, exercise or injury^33, 35^. Similar to our observations in the epidermis, inhibition of autophagy in myoblasts *in vitro* interferes with fusion [but not myoblast differentiation], but the mechanism of action for this has not been determined^33^.

Trophoblast differentiation represents another example for a potential correlation between autophagy and syncytium formation. The outer trophoblast cells in early mammalian embryos fuse with each other and form a multinucleated syncytiotrophoblast, which later establishes direct contact with the maternal blood that reaches the placental surface. Apart from enabling the exchange of nutrients, the syncytiotrophoblast also provides a barrier against maternal-fetal transmission of pathogens. Autophagy induces in both trophoblast layers, but the highest level of autophagy has been documented in the syncytiotrophoblast during cell fusion^36, 37^. Reduction of autophagy by Atg16L1 knockdown [even after rapamycin treatment] in both human and mouse placenta increases bacterial colonialization, but again it is not clear by what mechanism^36, 38^. Extrapolating from our own results, it is possible that also in this system, autophagy and syncytium formation are connected, in that autophagy leads to syncytilisation and thereby establishes the barrier function of the syncytiotrophoblast.

This leads us to speculate that perhaps in the *Drosophila* epidermis too, syncytium formation during wound healing may serve a protective function. We have seen that autophagy and syncytium formation are not necessary for wound closure as such, but that autophagy induces events indicative of enhanced barrier formation such as elevated levels of integrin in the apical compartment, a thicker basal lamina and a thicker outer cuticular region^39^. This may reinforce the freshly closed wound and protect it better against the external pathogens, especially in cases where the cuticle was compromised.

Another function of autophagy in the cells surrounding the wound may be to clear up debris. This is also seen in other instances. Autophagy is induced during Planarian regeneration in healthy cells adjacent to the wound and dead cells are phagocytosed by them^10, 40^. The cells surrounding the wounds in our experiments, also engulfed the wound debris^6, 13^. Whether multinucleated cells phagocytose and digest debris more efficiently^41^ remains to be seen.

Atg1/Ulk can activate myosin and drive autophagosome trafficking both in *Drosophila* and in mammalian cell lines^30^. This may explain the appearance of large cortical and cytosolic actomyosin filaments when autophagy was deregulated. It was perhaps counterintuitive that when the syncytium was wounded, the rate of the formation of the actin cable around the wound and subsequent wound closure were reduced. However, if both autophagy and wound closure need myosin activation, it is conceivable that autophagy might compete for regulators of actin assembly or contractility and thereby slow down actomyosin cable formation and wound closure. We saw that activation of actomyosin is not the cause but the consequence of the lateral membrane disassembly so that plasma membrane defects and wound healing delay are completely improved when autophagy reduced.

It is becoming clear that beside organelles such as ER, Golgi or mitochondria also the plasma membrane can supply the lipids or membrane required for autophagosome formation^2, 3^. While we see the large reservoir of lateral membrane being depleted in the epidermis, we do not know whether the material that is removed gives rise to autophagosomal membrane or reaches this compartment as cargo. Some of the components acting in the early steps of autophagosome assembly such as Atg16 and Atg9 are localized at the plasma membrane under normal conditions, where they directly interact with junctional components such as connexin (component of gap junction) and Patj (component of the apical cell polarity complex); they act in the delivery of plasma membrane or lipids to autophagosomes and in a polarized cell could contribute to the identification of the membrane domain to be used for autophagosome assembly^3, 42–45^. Why mitochondria are spared in the epidermis, whereas all other organelle membranes are depleted, is not clear.

Finally, in a *Drosophila* pathophysiological model for nuclear laminopathies the amount of nuclear membrane and its folding are increased^23, 25, 27–29^. Over-activation of autophagy improves the morphology and integrity of the nuclear envelope caused by Lamin B or Kugelkern overexpression. In summary, fine tuning of autophagy activity during development and tissue homeostasis is important for plasma membrane integrity.

## Methods

### Fly stocks

Fly stocks and crosses were maintained at 25 °C under a 12:12 h light/dark cycle at constant 65% humidity on standard fly food. All stocks were in a *white* genetic background. We used the *A58-Gal4* driver line to express UAS-constructs in the epidermis^13^ from the end of the first instar onward. Fly genotypes are listed in **Table 1**, fly stocks in **Table 2**.

**Table 1.**
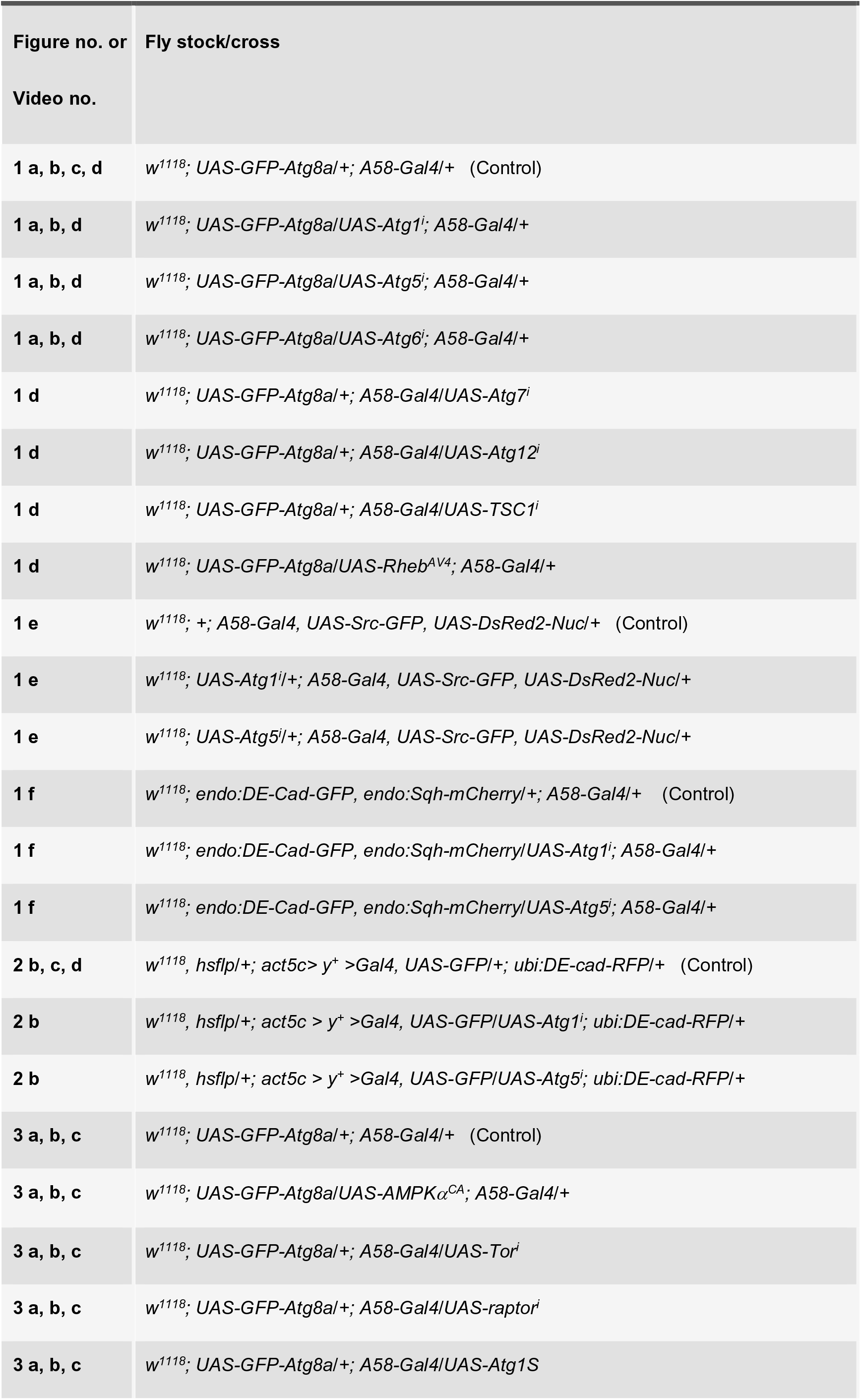
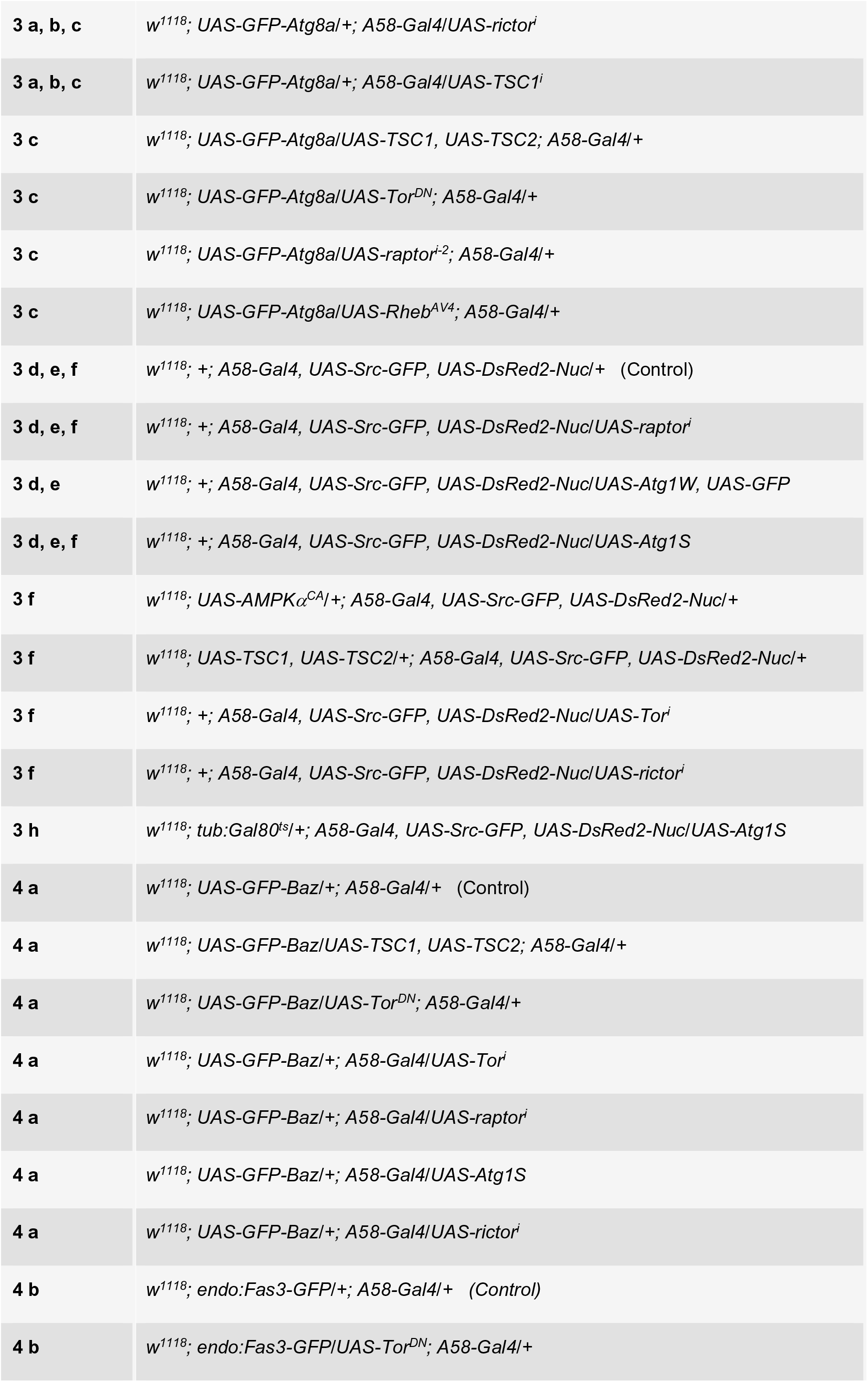
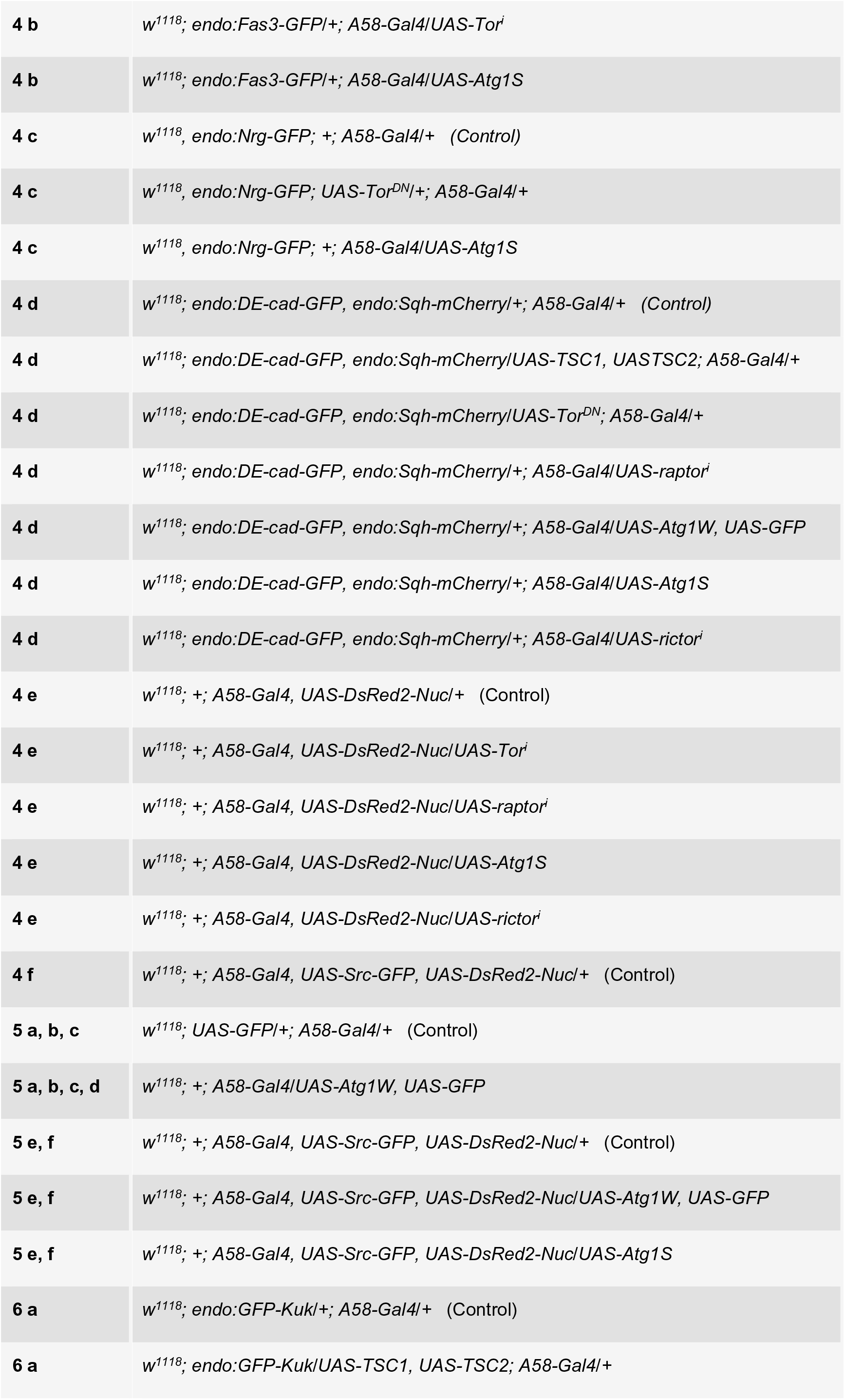
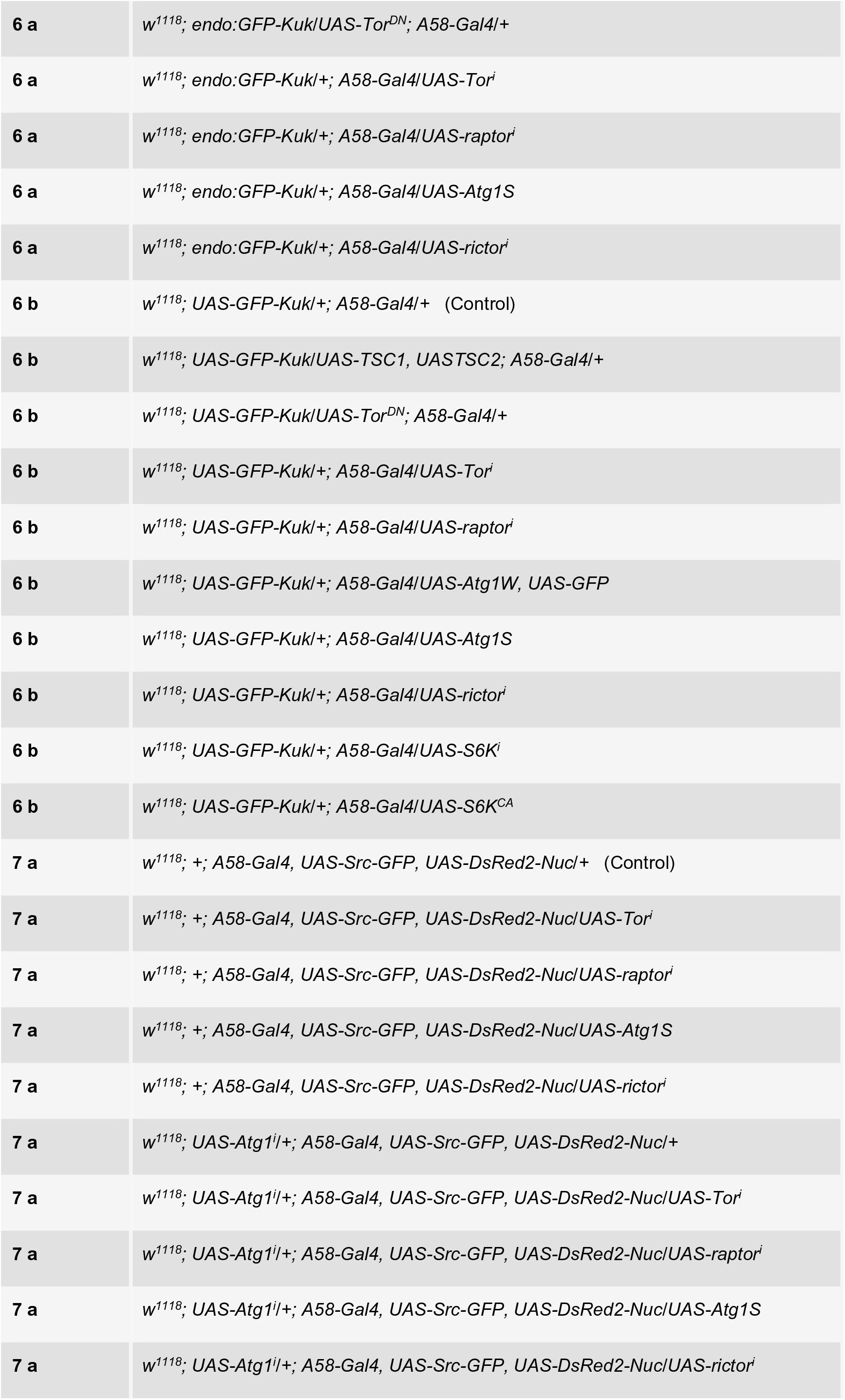
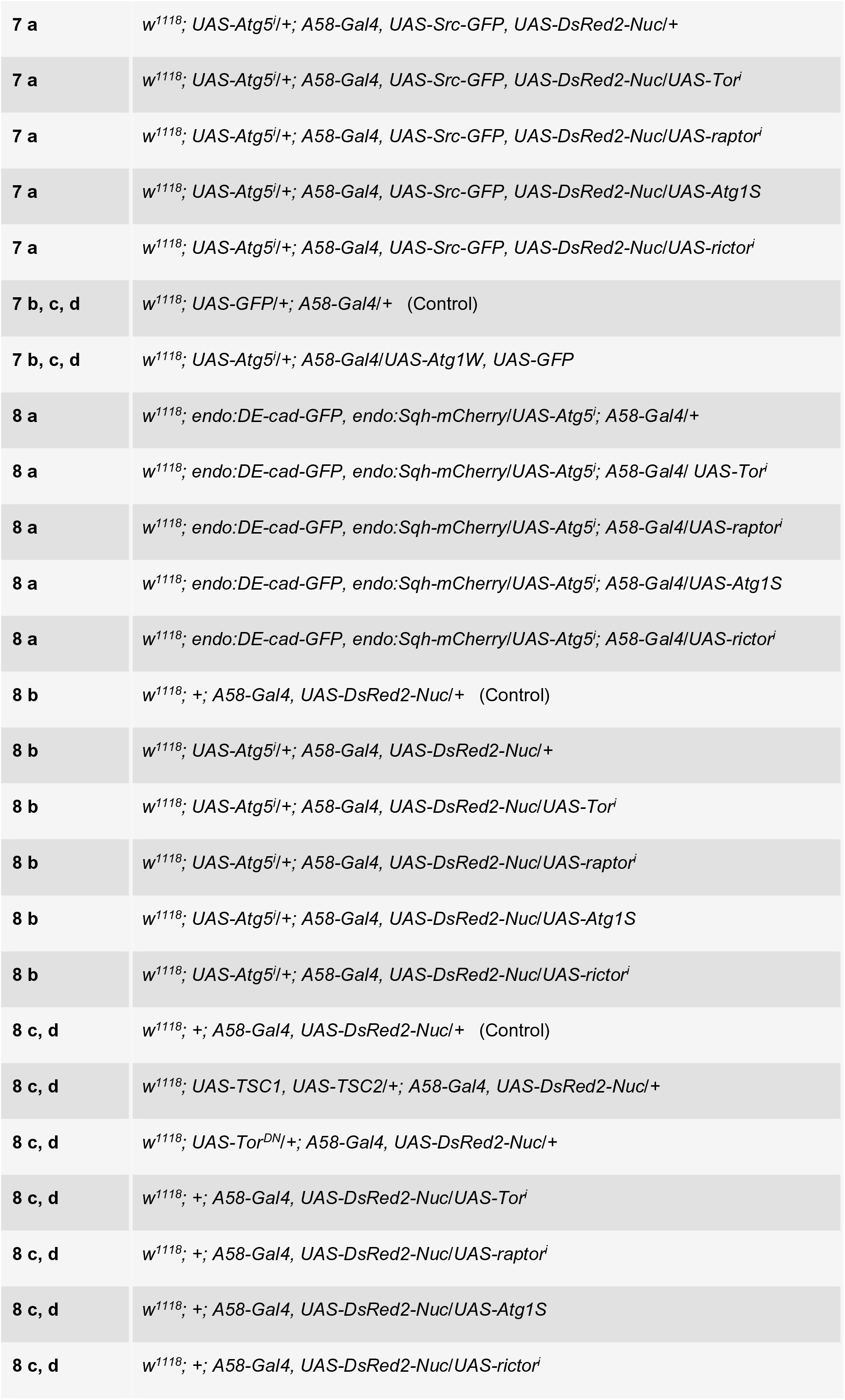
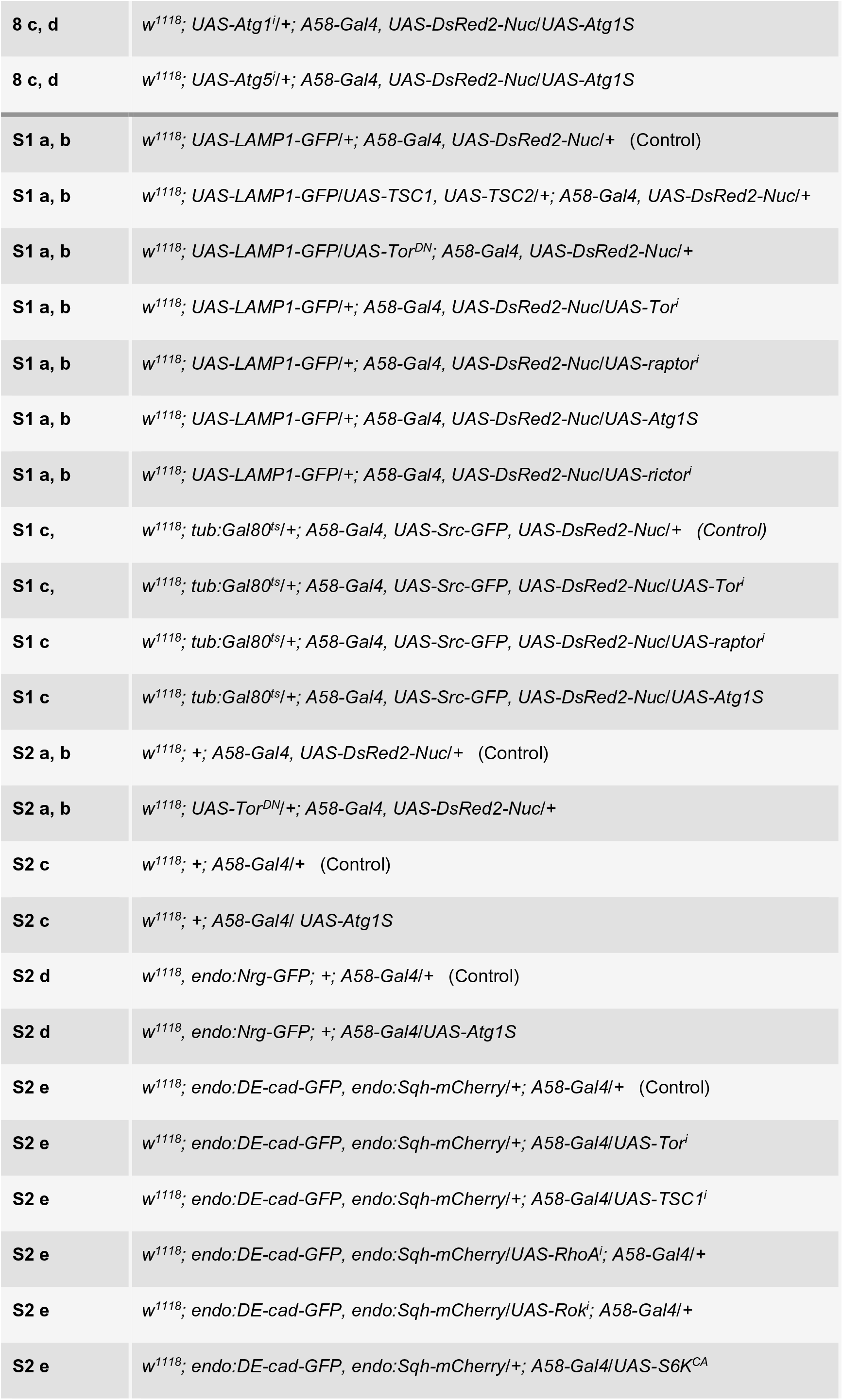
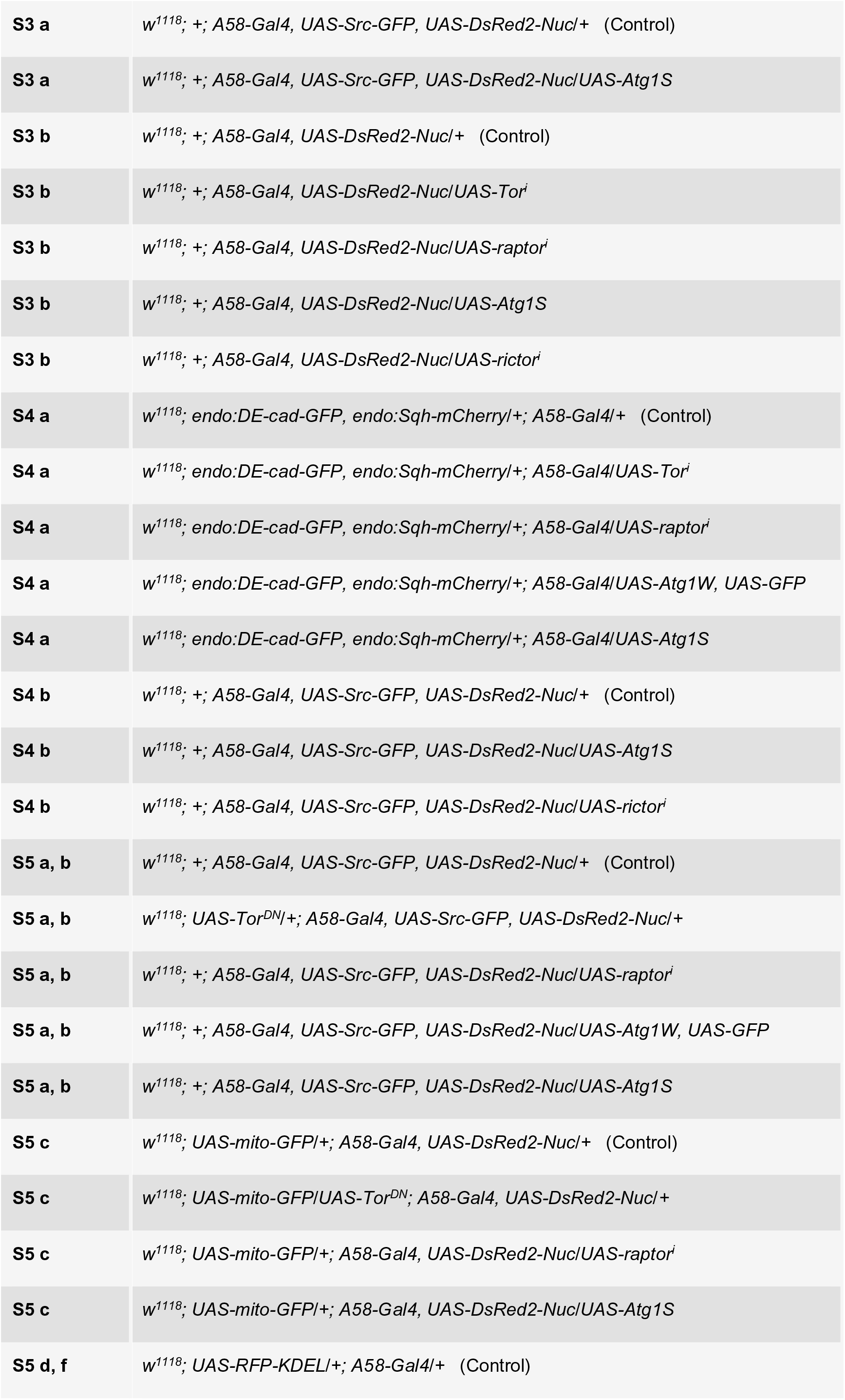
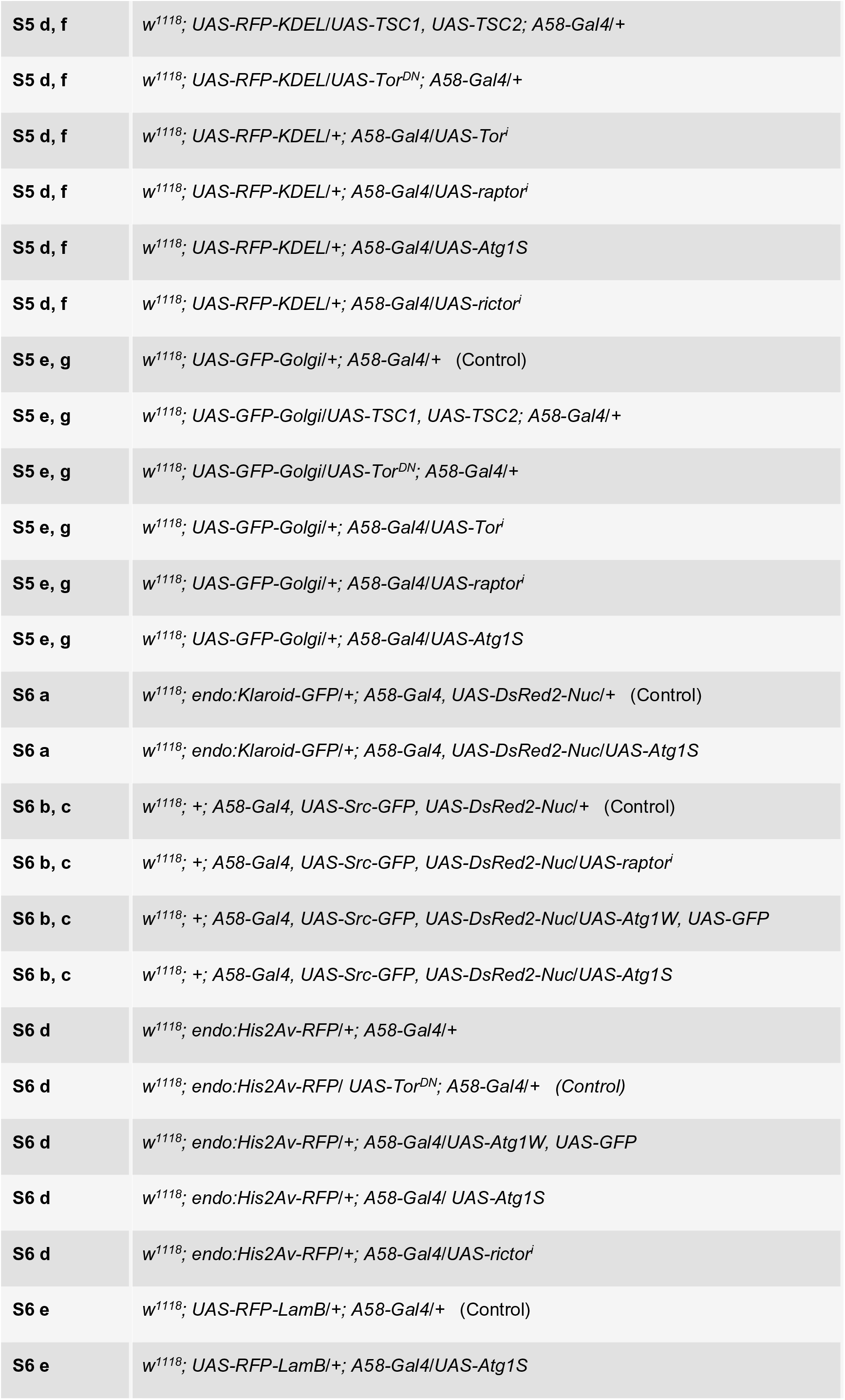
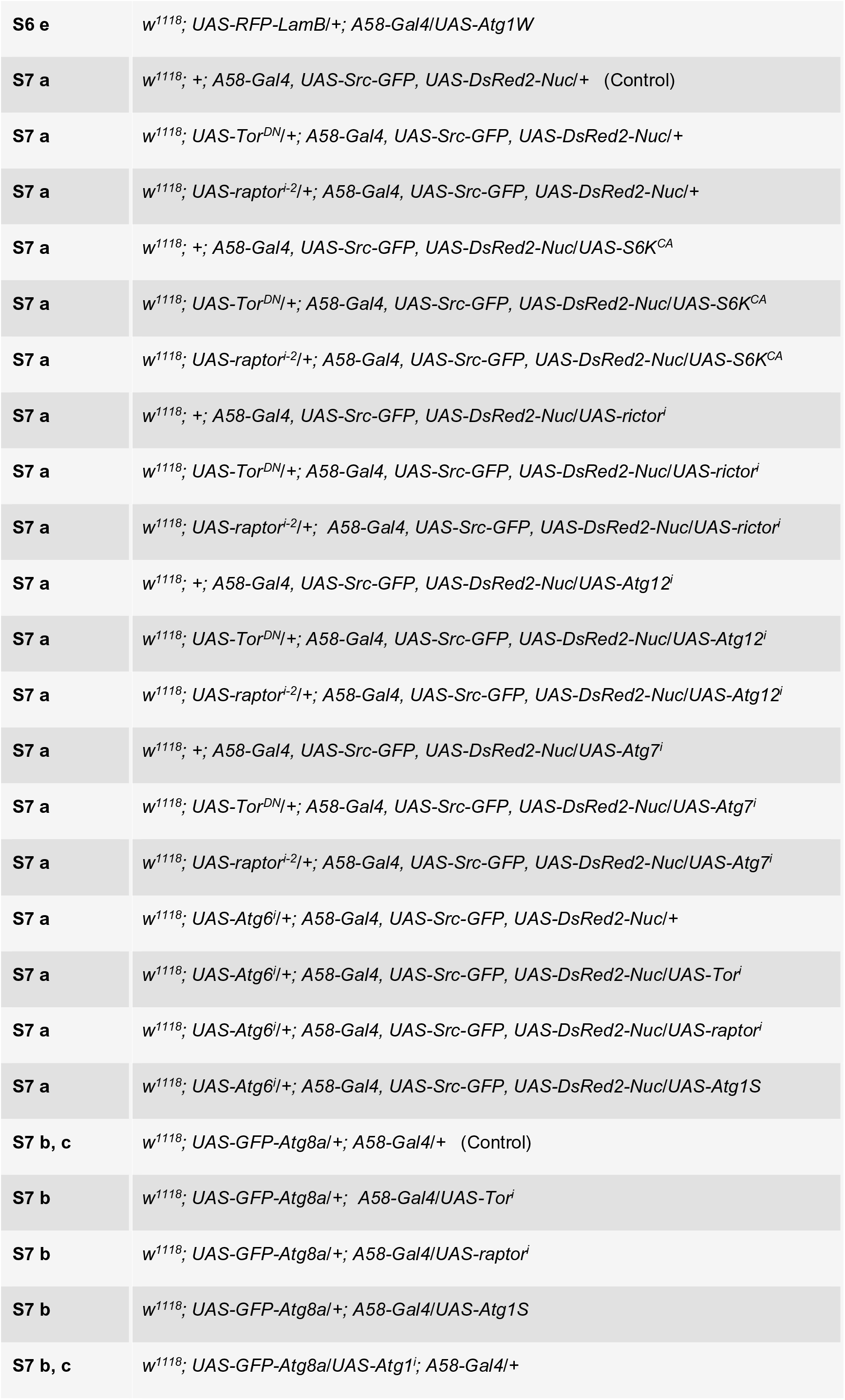
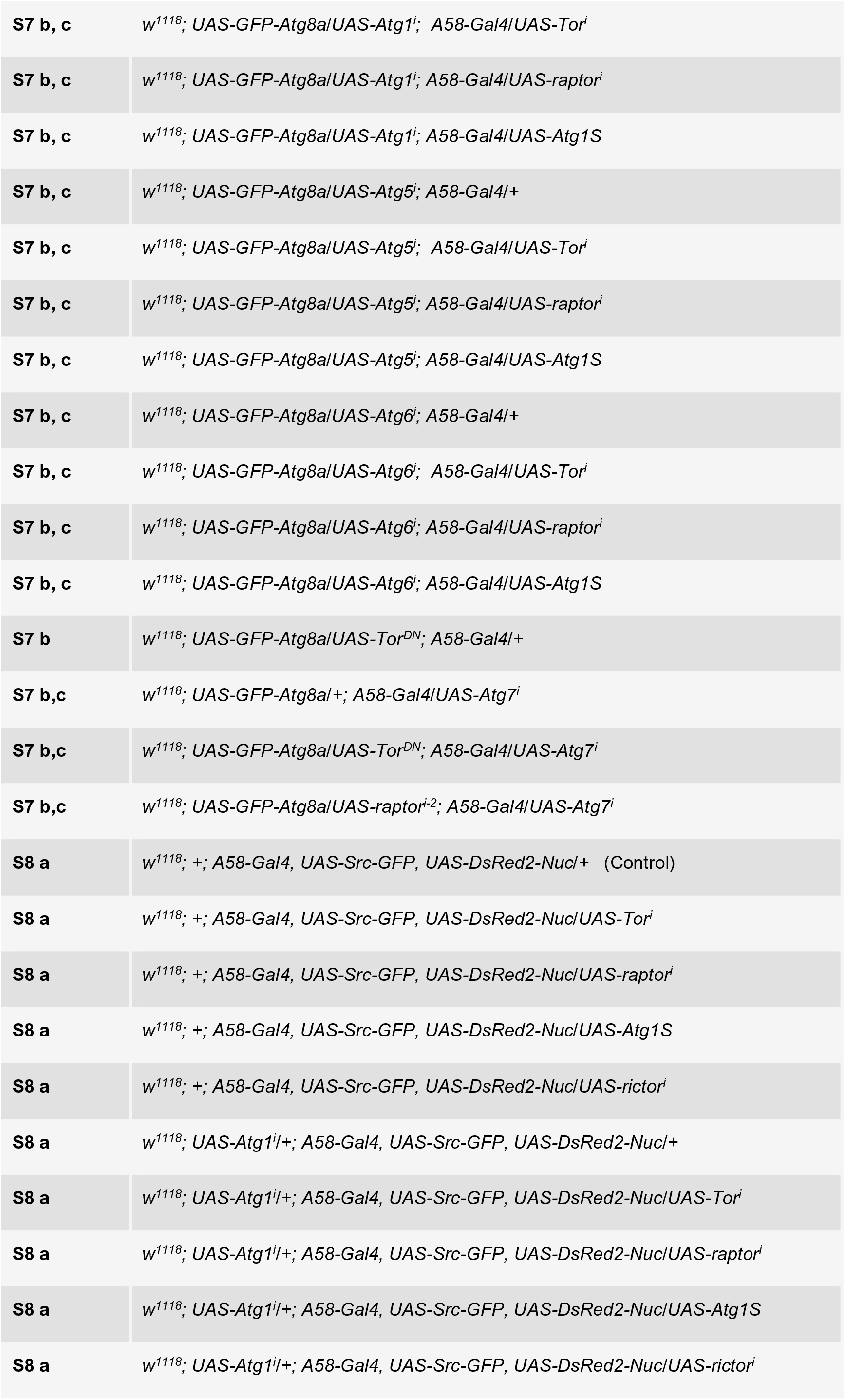
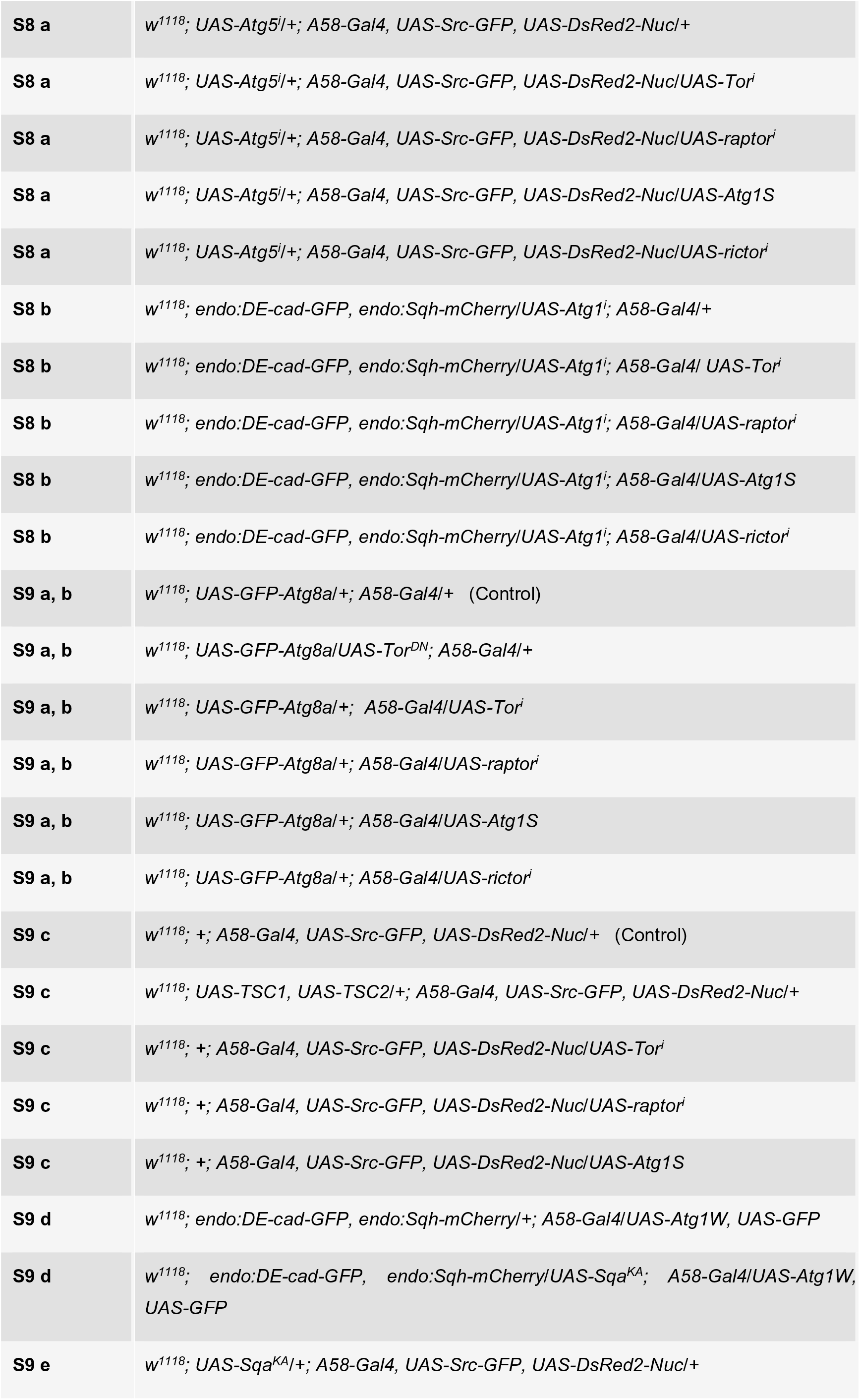
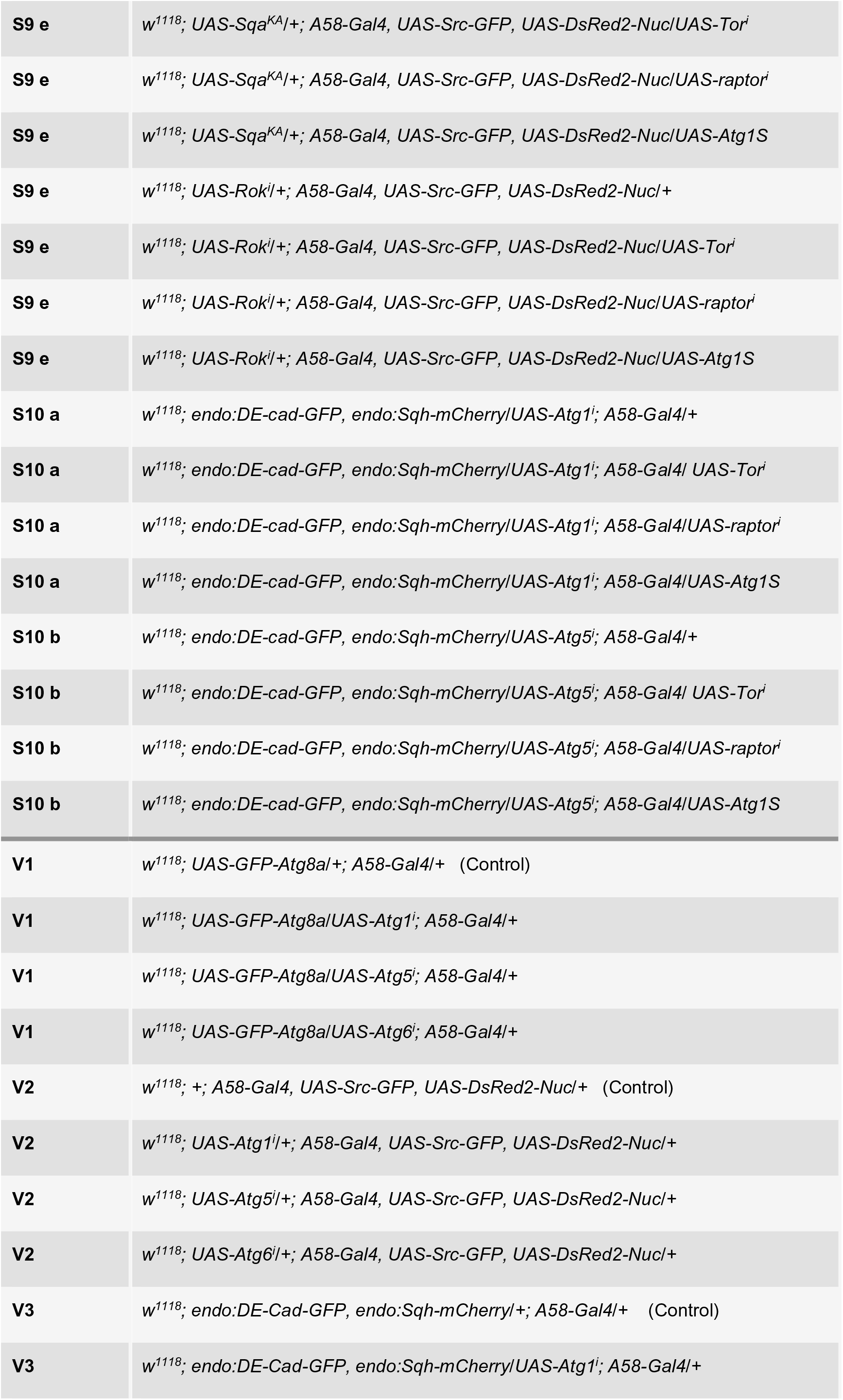
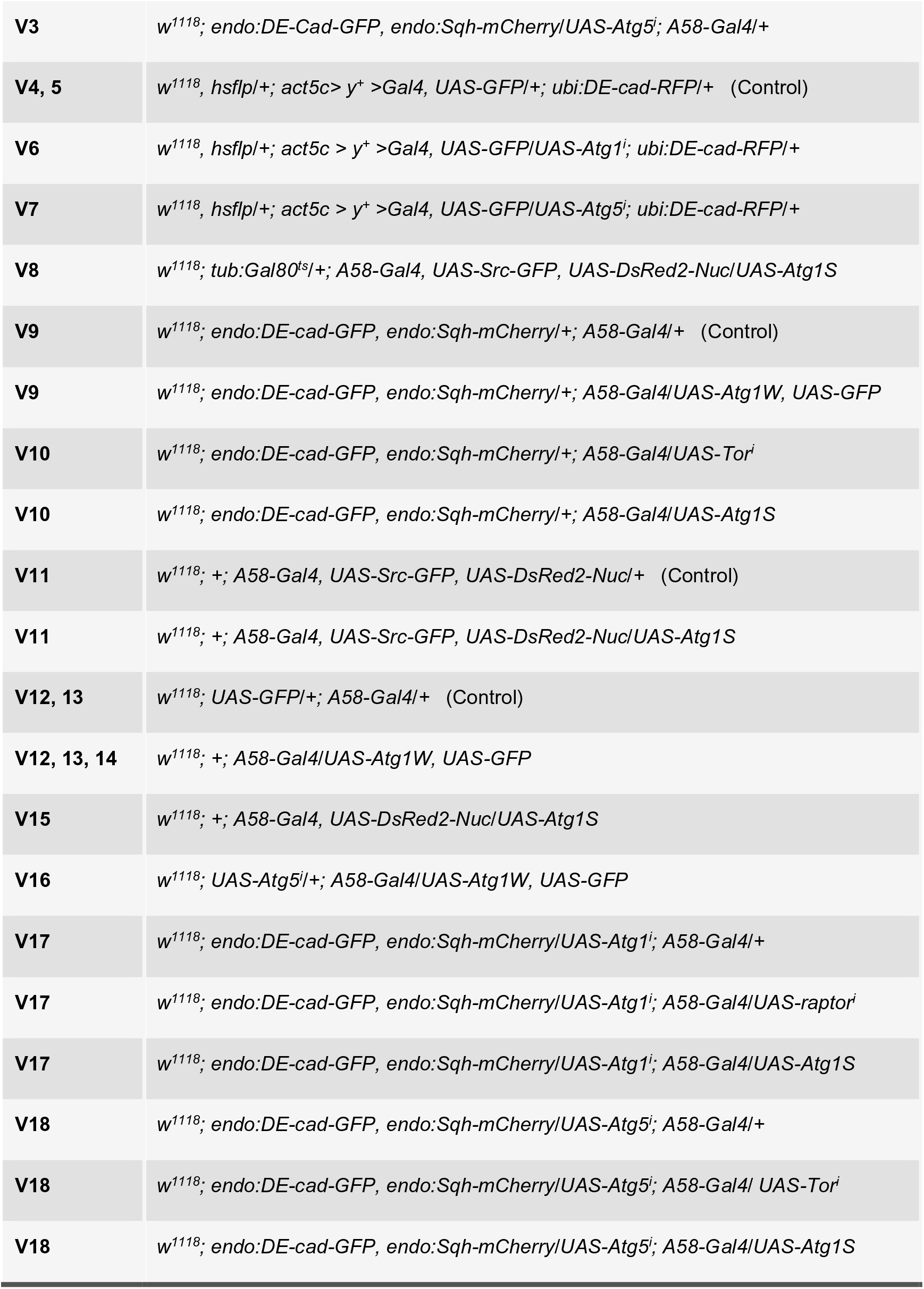
List of genotypes used in experiments.

**Table 2.**
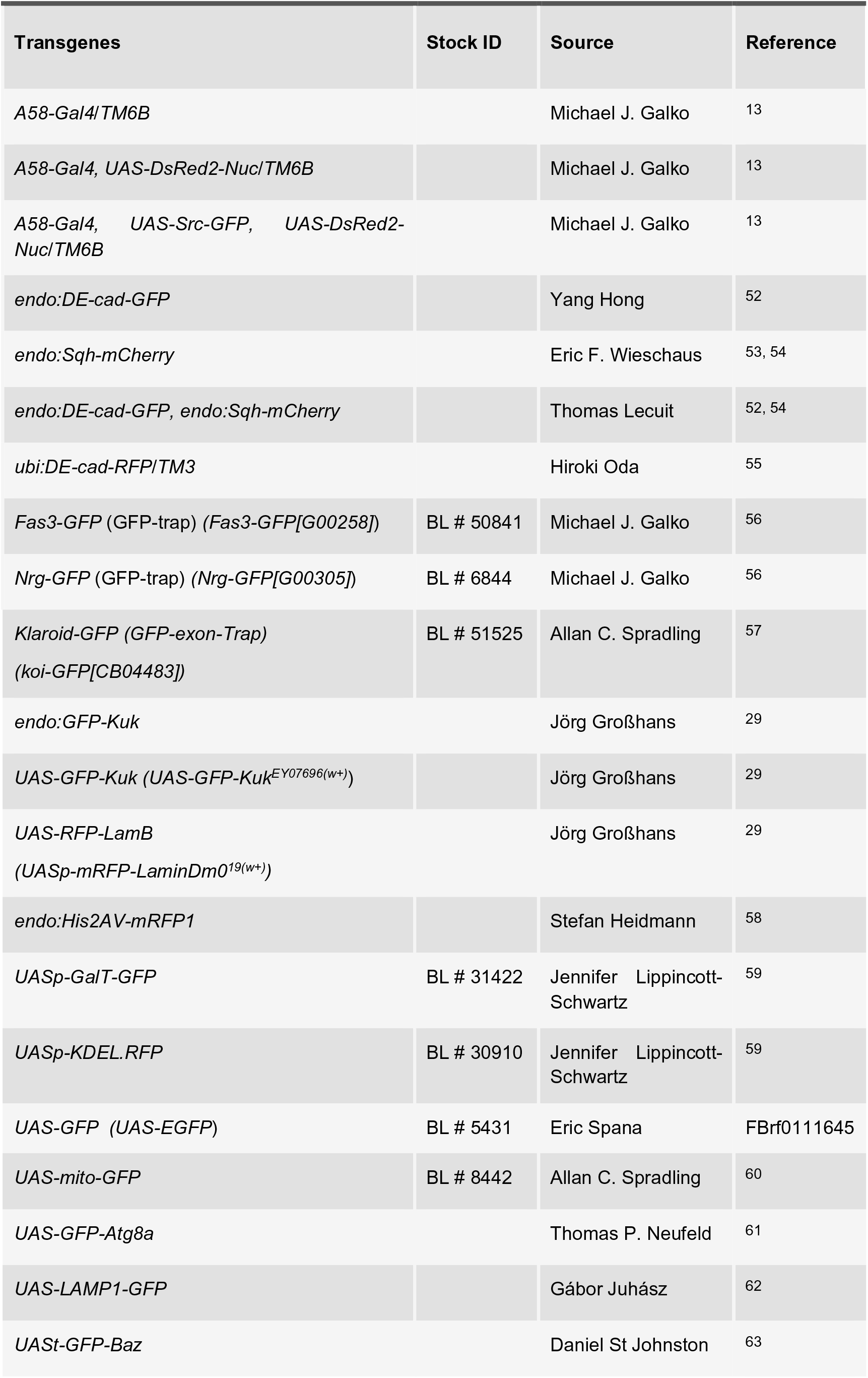
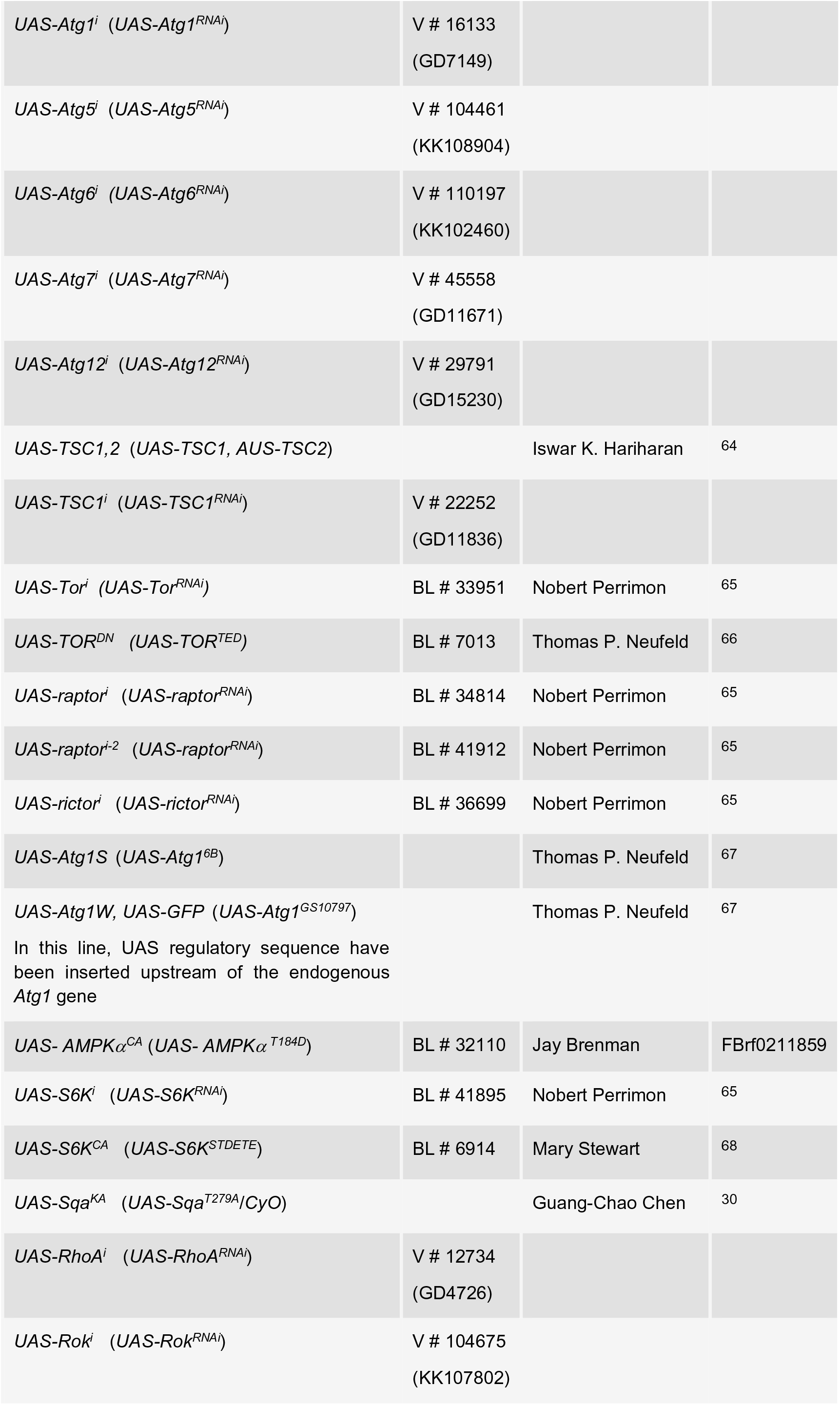
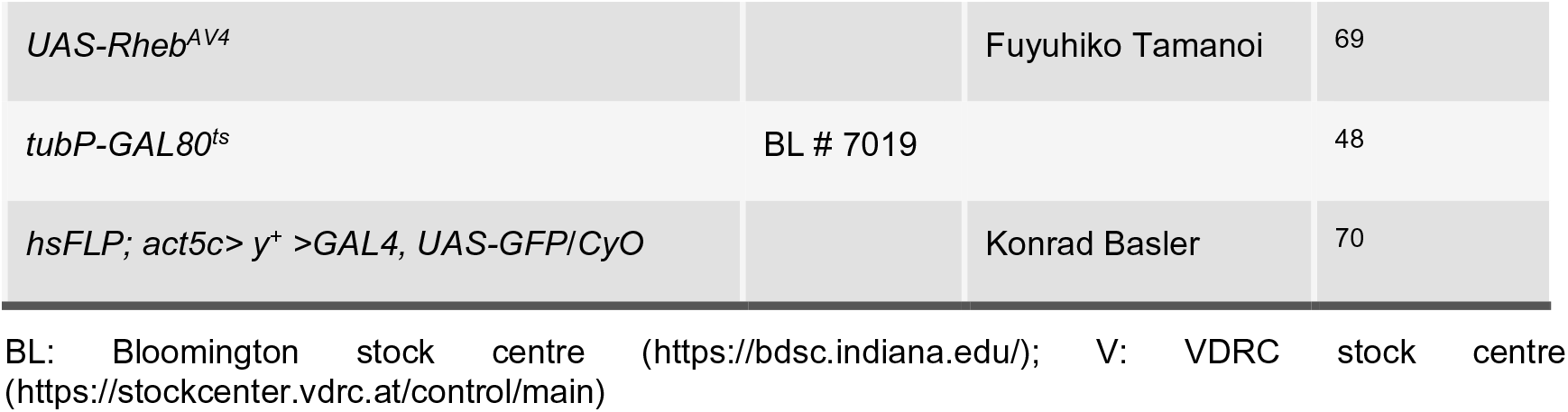
Fly stocks used in experiments.

### Flip-out experiments

*Atg1^i^*, *Atg5^i^, raptor^i^ and Atg1S* clones in the epidermis were generated with a flip-out system^46, 47^. Female flies with the genotype *w^1118^, hsflp; act> y^+^ >Gal4, UAS-GFP/*CyO; + were crossed with *w^1118^; UAS-Atg1^i^; ubi:DE-cad-RFP/TM6B or w^1118^; UAS-Atg1^i^; ubi:DE-cad- RFP/TM6B* or *w^1118^; endo:Sqh-mCherry; raptor^i^* or *w^1118^; endo:Sqh-mCherry; Atg1S* males or in control experiments with *w^1118^; +; ubi:DE-cad-RFP/TM3* or *w^1118^; endo:Sqh-mCherry; +* males. Clones were identified by GFP expression under the act5C promoter. Mating was carried out at 18 °C and at this condition no GFP was detected. The flip-out was induced in mid-stage L2 larvae by heat shock (15 or 30 min in a 37 °C w*ater bath*). The larvae were incubated for 30 h at 25 °C followed by wounding and imaging.

### Gal80^ts^ experiment

The temperature-sensitive Gal80 system (Gal80^ts^) was used for time-controlled induction of UAS-constructs^48^. *w^1118^; tub:Gal80^ts^; A58-Gal4, UAS-Src-GFP, UAS-DsRed2-Nuc/TM6B* females were crossed to males carrying the target UAS constructs at 18 °C; at this temperature no *Src-GFP* was detected. For inactivation of Gal80^ts^, larvae were heat shock in a 30 °C w*ater bath* for 6, 18 or 24 h. Less than 5 h heat shock was not sufficient to induce the *Src-GFP* marker in the epidermis. Live imagining of larvae with Gal80^ts^ was carried out at 30 °C.

### Chloroquine treatment

Mid-L2 Larvae were transferred to fresh medium containing 3 mg/ml chloroquine (Sigma-Aldrich Cat. # C6628) and 0.3% Erioglaucine disodium (Sigma-Aldrich Cat. # 861146; food colouring to monitor food uptake) for 14 h before imaging.

### Immunostaining

For immunofluorescent staining, L3 larvae were dissected in phosphate buffered saline (PBS), fixed in 4% formaldehyde or 4% paraformaldehyde for 30 min at room temperature (RT) and blocked in 1% BSA and 0.3% Triton-X for 2 h at RT. Primary and secondary antibodies were incubated for 2 h at RT. The tissues were mounted with Fluoromount-G for confocal imaging. Antiserum dilutions and sources: Bazooka (1:500, from A. Wodarz), a-PKC (1:500, Santa Cruze, ID-Nr #406), Fas3 (1:2000, DSHB, #7G10), βPS-integrin (1:200, DSHB #CF.6G11), anti-rabbit or anti-mouse immunoglobulin-G labelled with Alexa 488 (1:200; Invitrogen). BodiPy 493/503 (ThermoFisher Cat. #D3922) was used at 1: 500.

### FLIP procedures

A modified FLIP procedure^49^ was applied to the dorsal larval epidermis in the abdominal segments A3–A5 of L3 instar larvae on an inverted spinning disk microscope equipped with FRAPPA and a 40×/1.3 oil objective lens (see section **Microscope setup).** 7–19 frames were acquired before bleaching. 16-bit depth images were taken at a magnification of 0.163 μm/pixel acquiring a z-stack of 2.1 μm. Photobleaching (one bleaching event per cell, mid-cytosolic plane) was performed using 100% laser power over 3 or 20 minutes.

### Immobilizing larvae

Early L3 larvae were anaesthetised with diethyl ether^50^.

### Laser ablation

All laser ablations and wounding experiments were performed exactly as described previously (Kakanj et al. 2020). For single-cell wounds, the nucleus was targeted with 1 pulse/μm UV laser power of ∼0.30 μJ energy (measured on the objective lens). For laser cuts at the plane of the apical plasma membrane without inducing a wound healing response the UV laser was set to 1 pulse/μm of ∼0.25 μJ energy.

### Microscope setup and live Imaging

We used an inverted spinning disk confocal microscope (Nikon TiE model with Yokogawa, model no. CSU-X1 system) with a nano-positioning Piezo Z stage control system (NanoScan, model no. OP400, Prior Scientific), Plan-Fluor 40×/1.3 numerical aperture (NA) oil-immersion differential interference contrast (DIC) objective or Plan-Apochromat 60×/1.2 NA water-immersion objective; EMCCD camera (CamLink, model no. C9100-50; 1,000 × 1,000 pixels) controlled by Volocity v.6.3.57, with 488-nm and 561-nm channels. z-stacks were taken with a step size of 0.28 μm with 60×/1.2 and 0.24 μm with 40×/1.3 (45–95 z-stacks, covering a 10–30 μm depth) both for live and fixed specimens^50^. All figures and videos show merged z-stacks. After live imaging, larvae were returned individually into food vials and monitored for survival. We evaluated data only from larvae that survived at least two days post imaging. Live imaging used for the all images showing in this paper, except electron micrographs and images in **Figures 4e; 8b,c** and **Supplementary Figures 2a-c, 3c** which are from fixed specimens.

### Image processing

For image processing we used Volocity v.6.3.57 (PerkinElmer) or Fiji (National Institutes of Health).

### Image analysis

To quantify vesicles or puncta, the images were cropped to a uniform size and converted to 8-bit TIF files. Ilastik pixel-classifiers were trained separately using 6-15 images covering all genotypes for all markers. Separate classifiers were necessary since the nature of different ‘spots’ is different. The *Batch Processing* function in Ilastik was used to process all the images and *Simple Segmentation* results were exported. Custom Python script was then used to identify the ‘spots’ and to measure the number of spots in an area of 10000 µm^2^. The codes are available at https://github.com/sourabh-bhide/Analyze_Vesicles.

In the FLIP experiment, image intensities were measured in the bleached area and the sum of intensities in the area plotted over the time. All measurements were normalised to the pre-bleached value in the ROI.

### Data analysis

The plots were generated using the Python 3.7 library *Seaborn* (https://seaborn.pydata.org/generated/seaborn.boxplot.html). For statistical hypothesis testing, anindependent and non-parametric (Kruskal-Wallis) *t-tests* were performed for the mean number of ‘spots’ in control and experimental conditions in the graphs in Fig. 1d, 3c, 8d, S1b and S7c. We assumed unequal sample size and unequal variances and calculations were performed using the Scipy library from Python 3.7 and GraphPad Prism version 8.1. For all other graphs (Fig. S5f, S5g and S9b) ordinary one-way ANOVA statistical tests were performed after we confirmed normality using the Shapiro-Wilk test (alpha = 0.05). Values in this paper are presented as box plots. Box plot elements are: centre line, median; box limits, upper and lower quartiles; whiskers, 1.5x interquartile range; points, outliers. *P* values are indicated as follows: \**P* < 0.04; \*\**P* < 0.003; \*\*\**P* < 0.0002; \*\*\*\**P* < 0.0001 and lack of an asterisk or ns means non-significant (*P* > 0.123).

### Transmission electron microscopy

For transmission electron microscopy, early L3 *Drosophila* larvae were fixed and cryo-immobilized by high-pressure freezing as previously described^51^.

## Supporting information

Supplementary Video 1 | Autophagy during epidermal wound healing.

Supplementary Video 2 | Effect of autophagy on wound healing dynamics.

Supplementary Video 3 | Effect of autophagy on actomyosin cable formation.

Supplementary Video 4 | Syncytium formation during wound healing.

Supplementary Video 5 | Testing GFP de novo synthesis in response to wounding or laser cut.

Supplementary Video 6 | Effect of supressing Atg1 on syncytium formation during wound healing.

Supplementary Video 7 | Effect of supressing Atg5 on syncytium formation during wound healing.

Supplementary Video 8 | Temporal effect of autophagy induction.

Supplementary Video 9 | Effect of uncontrolled autophagy on actomyosin cable formation during wound healing.

Supplementary Video 10 | Effect of Tor signalling and uncontrolled autophagy on actomyosin cable formation.

Supplementary Video 11 | Effect of uncontrolled autophagy on wound healing.

Supplementary Video 12 | Effect of uncontrolled autophagy on lateral membrane integrity.

Supplementary Video 13 | Effect of uncontrolled autophagy on membranes.

Supplementary Video 14 | Uncontrolled autophagy and membrane leakage.

Supplementary Video 15 | Effect of uncontrolled autophagy on nuclei.

Supplementary Video 16 | Atg1 acts via Atg5.

Supplementary Video 17 | Block of Atg1 improves actomyosin cable and wound healing in uncontrolled autophagy.

Supplementary Video 18 | Block of Atg5 improves actomyosin cable and wound healing in uncontrolled autophagy.

## Acknowledgements

We are grateful to A.J. Garcia-Saéz, F. Papagiannouli, M. Rembold, M. Graef, N.L. Kononenko and S. Roth for critical reading of the manuscript, comments and helpful discussions. We thank A. Wodarz, M.J. Galko, L. Partridge and M. Uhlirova for antibodies and fly lines. We thank A. Schauss, F. Babatz, P. Zentis and C. Jüngst from the CECAD imaging facility in Cologne (University of Cologne, Cluster of Excellence in Ageing Research) for technical support and the Bloomington, VDRC and DGGR stock centers for fly strains. This work was funded by CMMC (Projekt-Nr. 16-RP, Fond 2635/8025/01) and CECAD through a grant by the Deutsche Forschungsgemeinschaft (DFG, German Research Foundation) under Germanýs Excellence Strategy – EXC 2030 – 390661388, Gefördert durch die Deutsche Forschungsgemeinschaft (DFG) im Rahmen der Exzellenzstrategie des Bundes und der Länder - EXC 2030 – 390661388 to M.L.

## Contributions

P.K. conceived the project, designed the experiments, performed all experiments; and prepared the figures and tables. P.K. and B.M. performed TEM analysis and S.B. performed all image quantifications. P.K. and M.L. analysed and discussed the data and drafted the manuscript.

## Supplementary information

Supplementary information is available within the same PDF and as mp4 Video.

## Supplementary Figure legends

**Suppl. Fig. 1.**
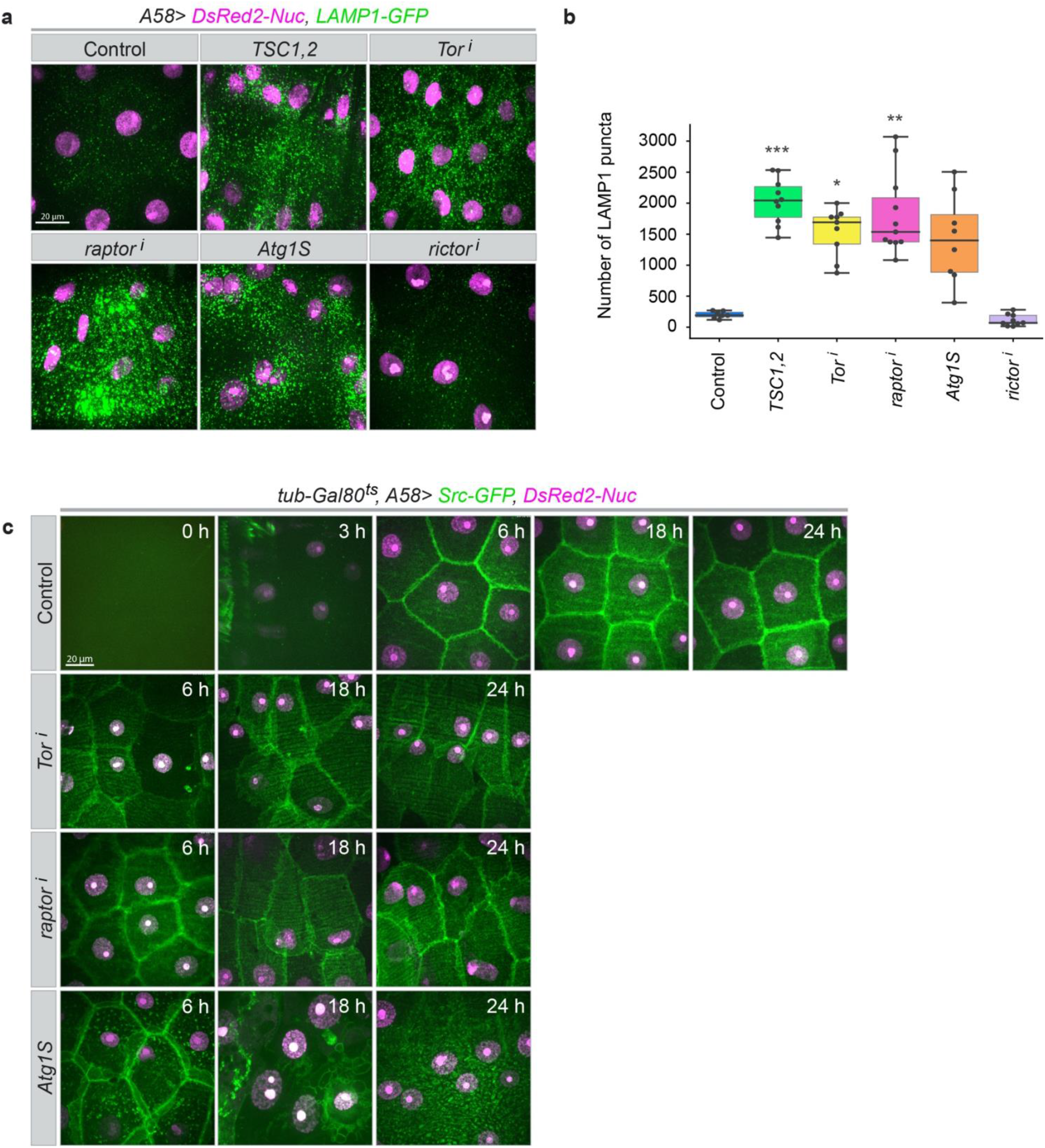
Effect of uncontrolled autophagy on epidermal organelles and membranes. **a,b,** Epidermis expressing markers for lysosomes (LAMP1, green) and nuclei (magenta) together with the indicated overexpression or RNAi constructs. **a,** Representative in images. **b,** Quantification of lysosomes (LAMP1) in an area of 10000 µm^2^. **c,** Temporally controlled activation of autophagy in larval epidermis expressing membrane (green) and nuclear (magenta) markers and the indicated constructs. Transgene expression was induced according to the time schedule shown in Fig. 3g. Each time point shown is a separate experiment, analysed at the indicated time post-activation The nuclear DsRed2 first becomes detectable after 3h, membrane outlines are seen clearly after 6h. Morphological defects are apparent at this time. **a,c,** z– projections of time-lapse series. **a,** n=15–30, **b,** n=7–11, **c,** n=15–30 larvae for each genotype for each time point. Scale bars, **a,c,** 20 μm. Genotypes of all images are listed in **Table 1**.

**Suppl. Fig. 2.**
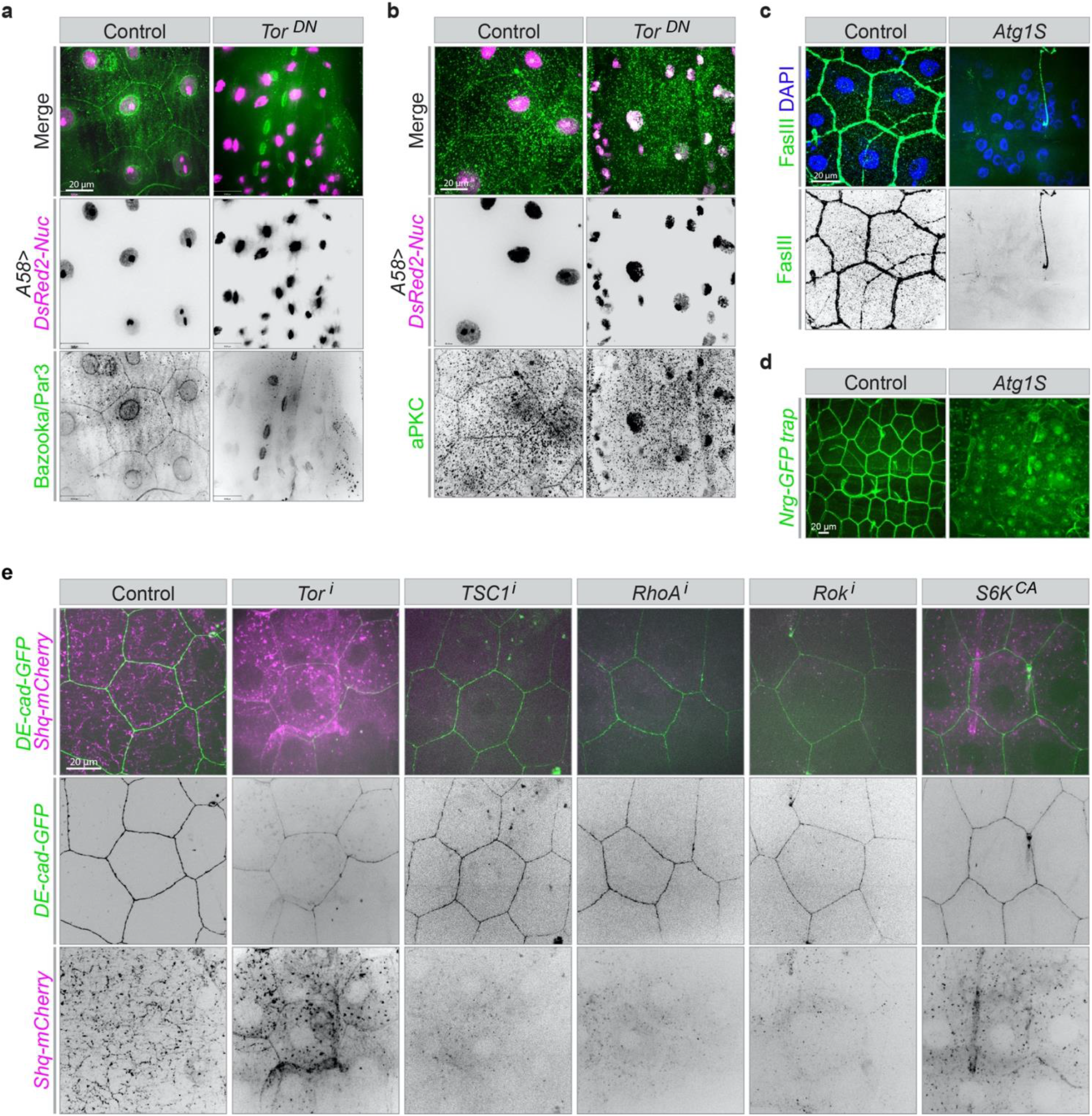
Effect of uncontrolled autophagy on cell polarity and junctions. **a-e,** Larval epidermis expressing the indicated markers and overexpression or RNAi constructs. **a-c,** Antibody stainings for junctional and polarity markers Bazooka/Par3 (**a**), aPKC (**b**) FasIII (**c**) on fixed epidermis. **d,e** In vivo imaging of larval epidermis expressing endogenously tagged, Nrg-GFP (**d**) or E-cadherin-GFP and Sqh-mCherry (**e**). Reduction of Bazooka/Par3, aPKC, FasIII, Nrg-GFP and E-cadherin-GFP by uncontrolled autophagy and reduction of actomyosin by TOR activation in TSC1i (compare to Fig. 4d). **a-e**, n=9–25 larvae each genotype, **a-e** z–projections. Scale bars, **a-e,** 20 μm.

**Suppl. Fig. 3.**
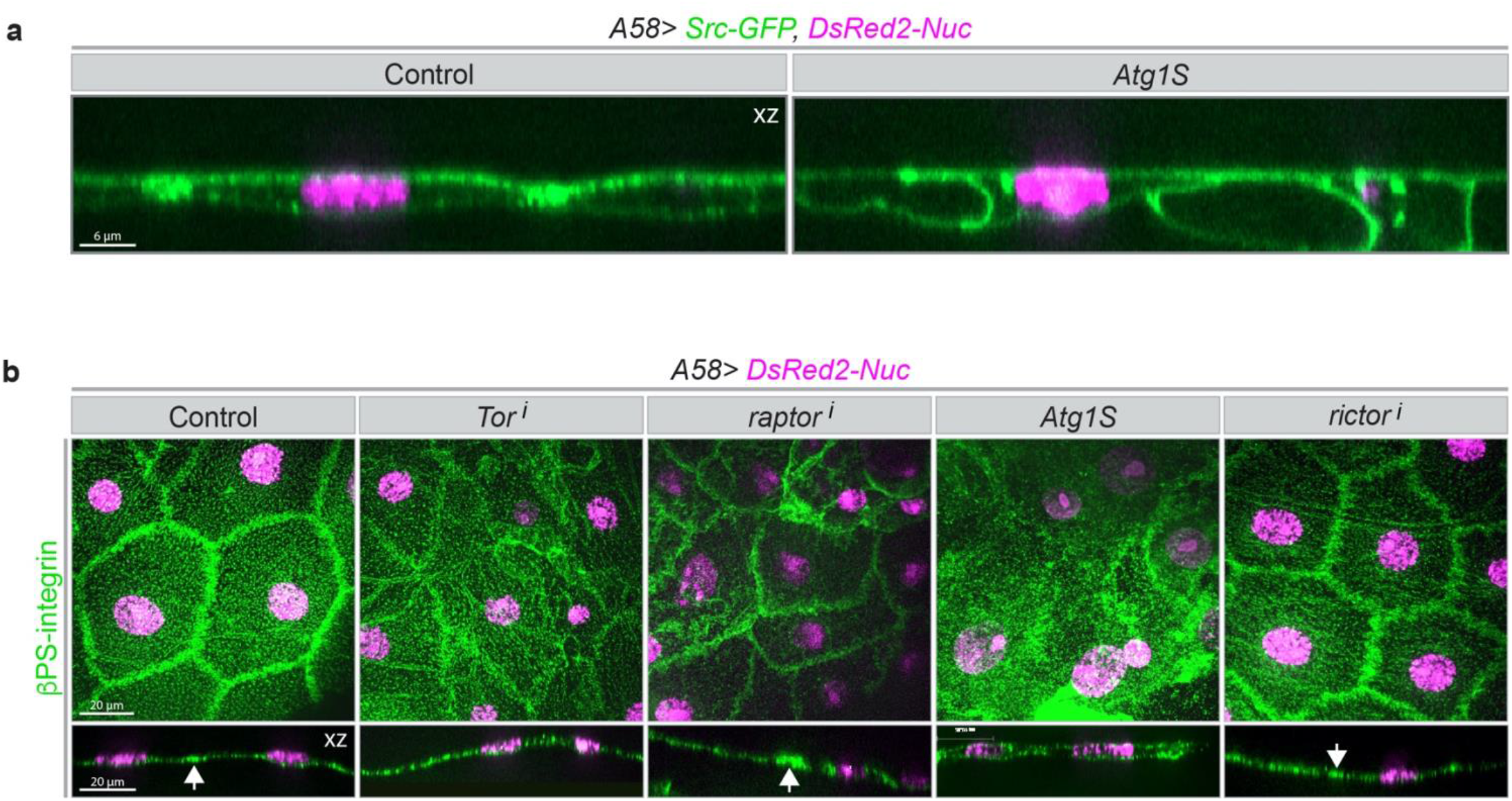
Effect of uncontrolled autophagy on actomyosin and integrin. **a-c,** Larval epidermis expressing the indicated markers and overexpression or RNAi constructs. **a,** Clonal analysis; Sqh-mCherry is expressed constitutively throughout the epidermis, whereas free GFP is induced in random clones under the control of *actin5c- Gal4* (see scheme in Fig. 2a). Excessive actomyosin stress fibres and cables are restricted to the clonal cells. **b,** Higher magnification z-section. **c,** Antibody staining forβ-integrin on fixed epidermis. Surface views and z-sections through the same samples. Arrows in the z-sections point to high accumulation of integrin along the folded lateral membranes. The z-section of the epithelium expressing Atg1S has integrin both in the apical and the basal membrane. **a,c**, z–projections, **a,b,** In vivo imaging. **a-c,** n=10–23 each genotype. Scale bars, **a,** 6 μm and **b,** 20 µm.

**Suppl. Fig. 4.**
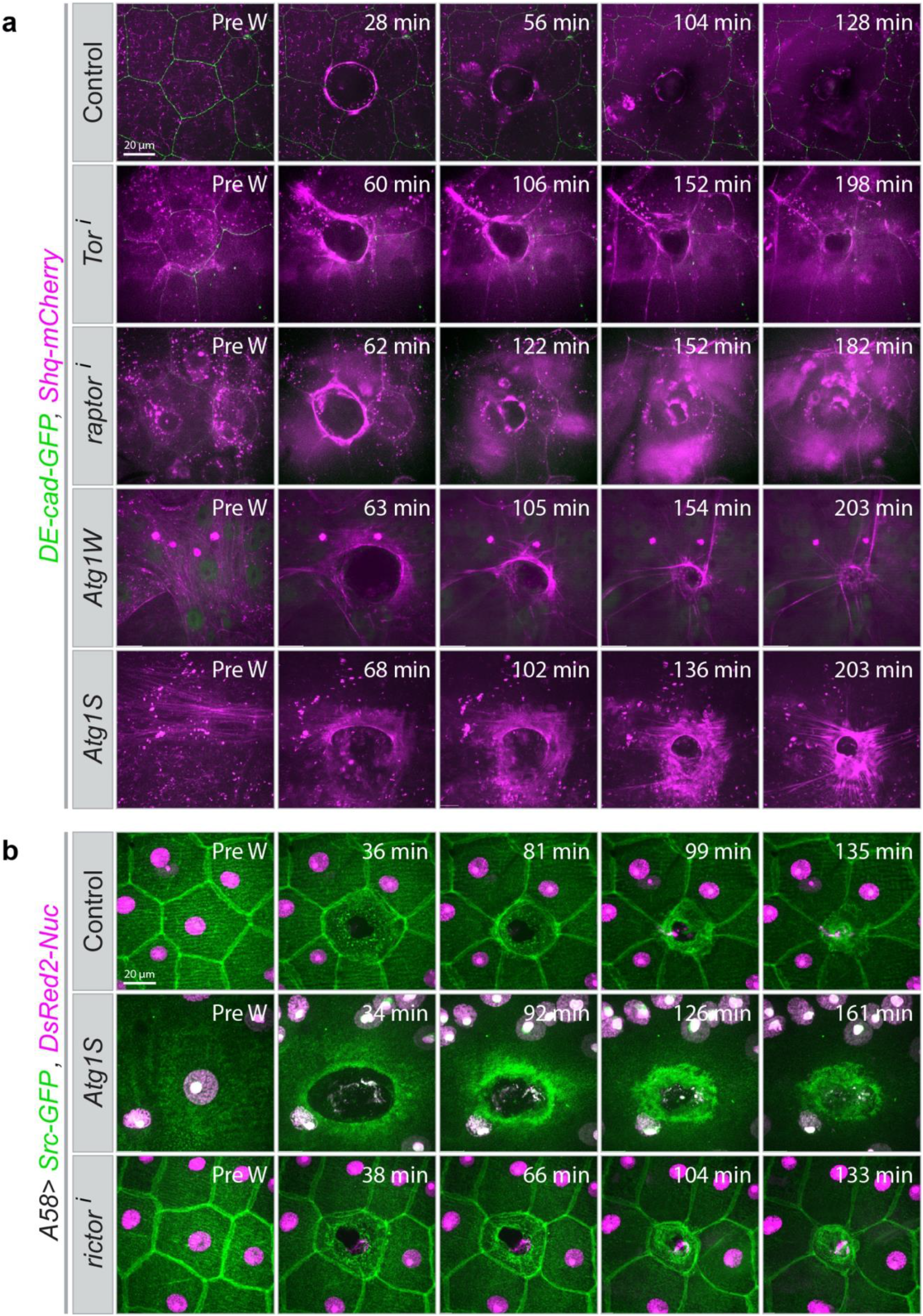
Effect of uncontrolled autophagy on wound closure. **a,b,** Larval epidermis expressing the indicated markers and overexpression or RNAi constructs before and after wounding**. a,** Uncontrolled autophagy leads to delayed but excessive actomyosin cable formation around the wound**. b,** Wound healing is delayed but completed. **a,b,** z–projections of time-lapse series, n=20–30 larvae each genotype. Scale bars, **a,b,** 20 μm. Images from **Supplementary Videos 9-11**.

**Suppl. Fig. 5.**
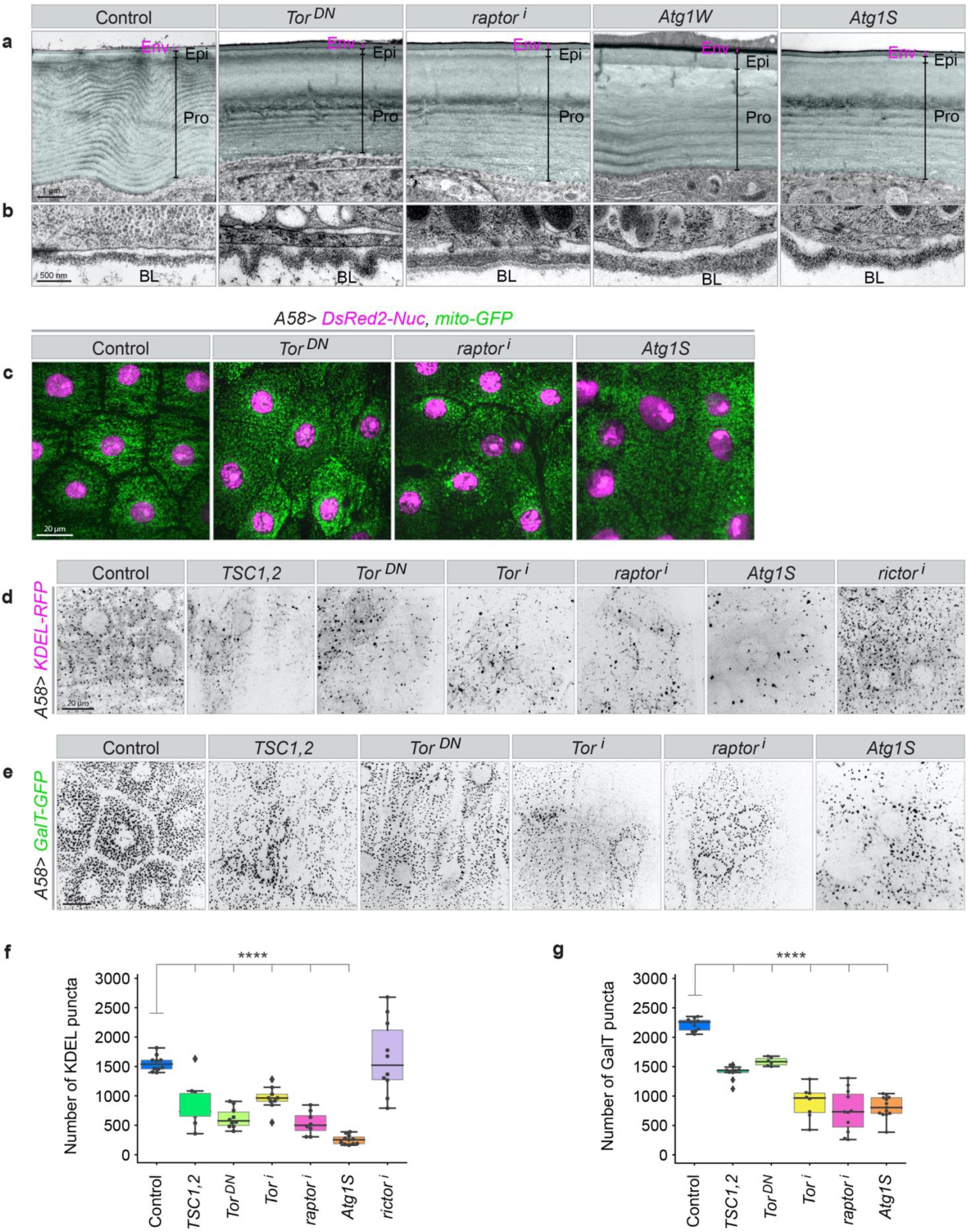
Effect of uncontrolled autophagy on epidermal morphology and organelles. **a,b**, Electron micrographs of the cuticle (**a**) and basal lamina (**b**) of larval epidermis expressing the indicated transgenes. Cuticle false-coloured in blue. Env: outermost envelope; Epi: epicuticle; Pro: procuticle, BL: basal lamina. **c-e,** Larval epidermis expressing markers for mitochondria (**c**), ER (**d**) and Golgi (**e**) together with the indicated overexpression or RNAi constructs. Mitochondria are not affected, ER and Golgi are reduced, as quantified in (**f**,**g**)**. f,g** number of ER and Golgi spots in 10000 µm^2^ epidermis in the indicated conditions. **c-e,** z–projections of in vivo images, n=20–40 larvae each genotype. Scale bars, **a,** 1 µm; **b,** 500 nm; **c-e,** 20 μm.

**Suppl. Fig. 6.**
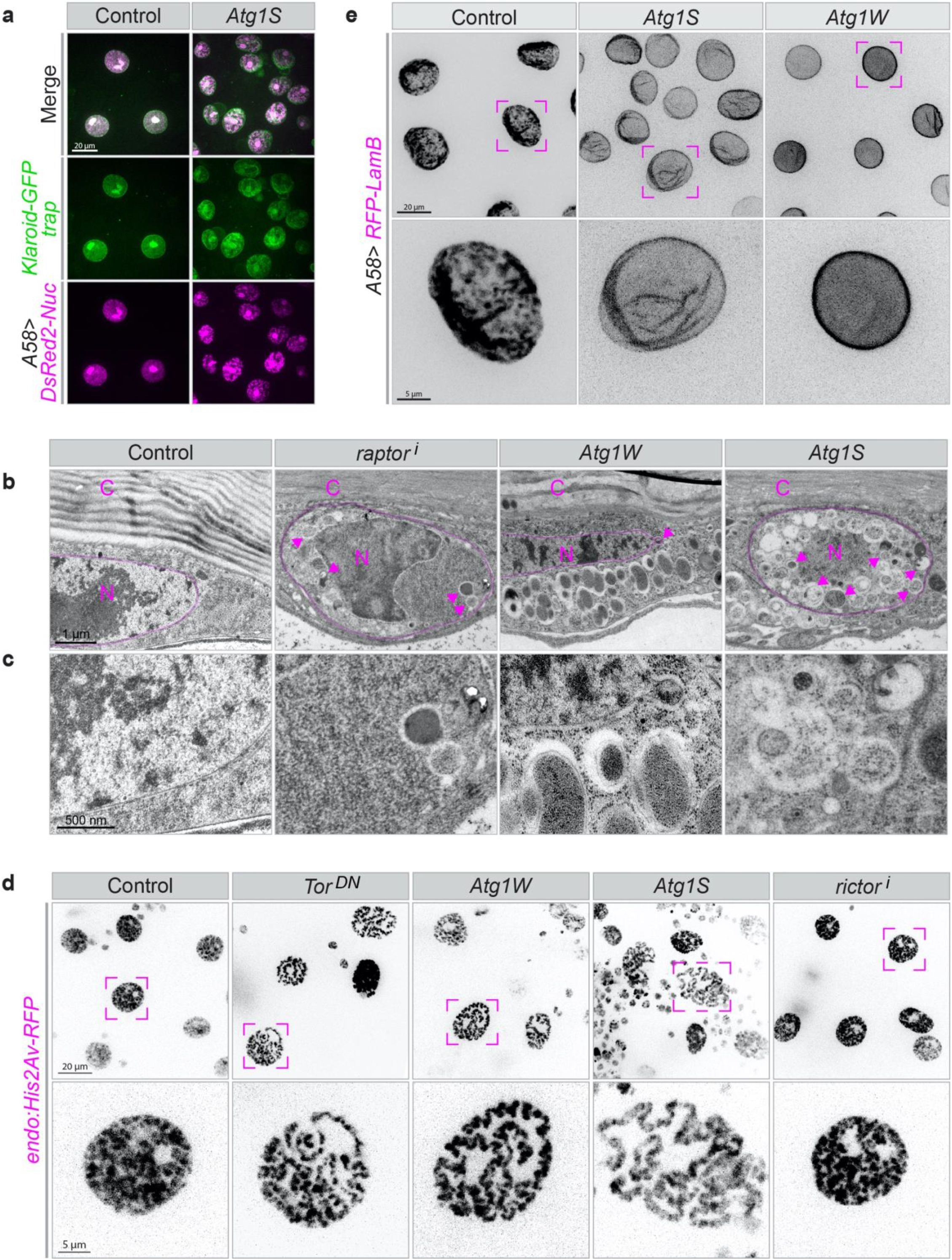
Effect of uncontrolled autophagy on nuclear morphology. **a,d,e,** Fluorescence and **b,c,** electron micrographs of larval epidermis expressing the indicated markers and overexpression or RNAi constructs. **a,** Klaroid-GFP is a GFP-insertion in the endogenous *klaroid* gene. **b,c**, Electron micrographs of cross sections of the epidermis. C: cuticle; N: nucleus; magenta arrows: autophagosomes. **c,** Details at higher magnification. **d,** Chromosome arrangement in epidermal nuclei after upregulation of autophagy. **e,** High levels of LamB induce lobulation and other nuclear defects, which are ameliorated if autophagy upregulated. **d,e,** Lower rows: higher magnification of the nuclei marked above. n=20–40 larvae each genotype. Scale bars: **a** 20 μm; **b,** 1 µm; **c,** 500 nm; **d,e,** upper row 20 µm; **d,e,** lower row, 5 μm. **Supplementary Video 15**.

**Suppl. Fig. 7.**
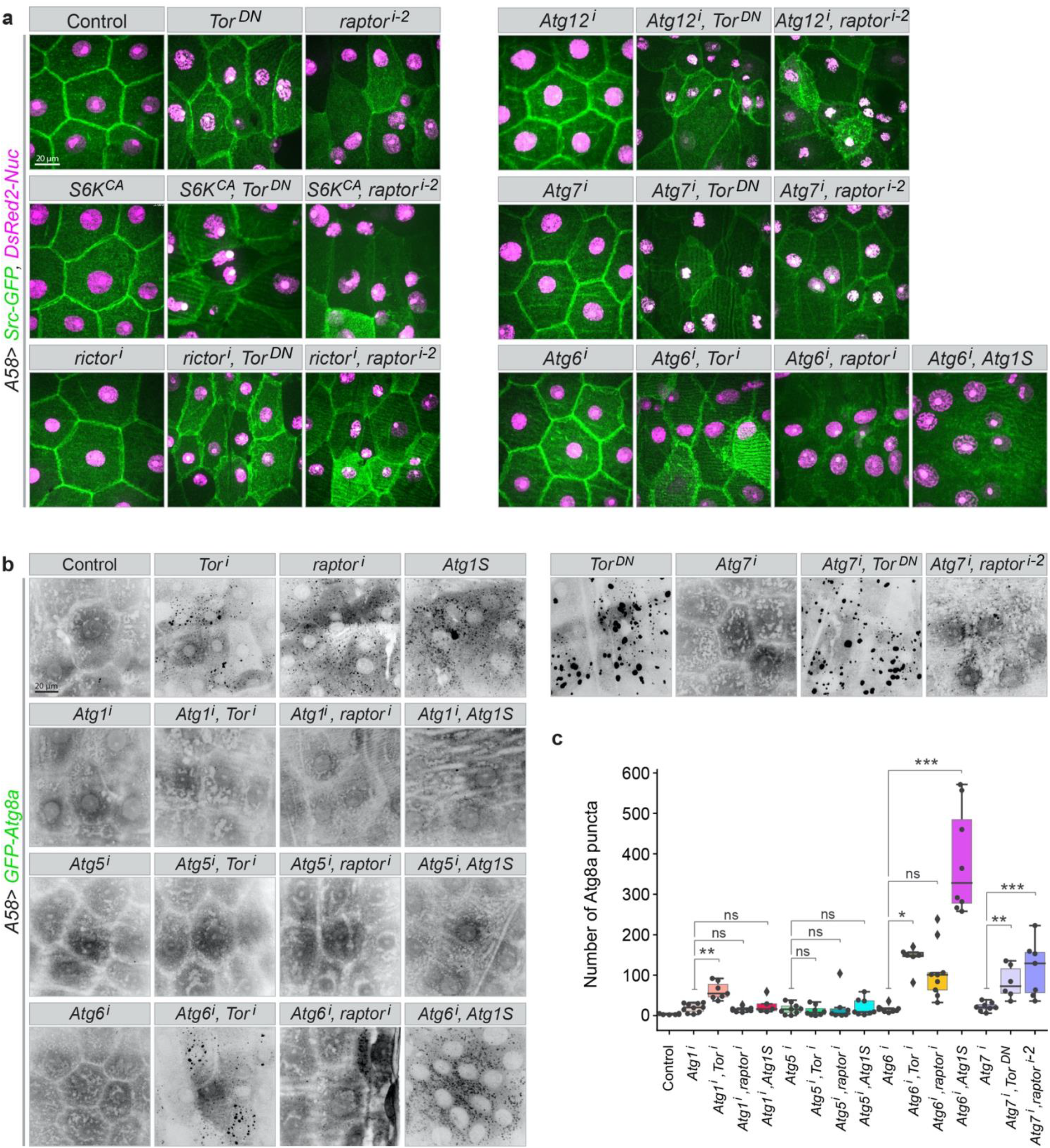
Epistasis of components in the TOR/autophagy pathway in epidermal morphology. Larval epidermis expressing the indicated markers and overexpression or RNAi constructs. **a**, The effect of reduced TOR signalling cannot be suppressed by inhibiting rictor, Atg6, Atg7, Atg12 or by constitutively active S6K. Note that the Atg6, Atg7 or Atg12 constructs do not suppress autophagy efficiently (see **Fig. 1d** and **Suppl. Fig. 7b,c**). **b,** Ability of RNAi constructs for components of the autophagy pathway to inhibit uncontrolled autophagosome formation. **c,** Quantification of autophagosomes in an epidermal area of 10000 µm^2^ in the indicated conditions; n=7–10 larvae each genotype. **a,b,** n=15–40 larvae each genotype. Scale bars: **a,b,** 20 μm.

**Suppl. Fig. 8.**
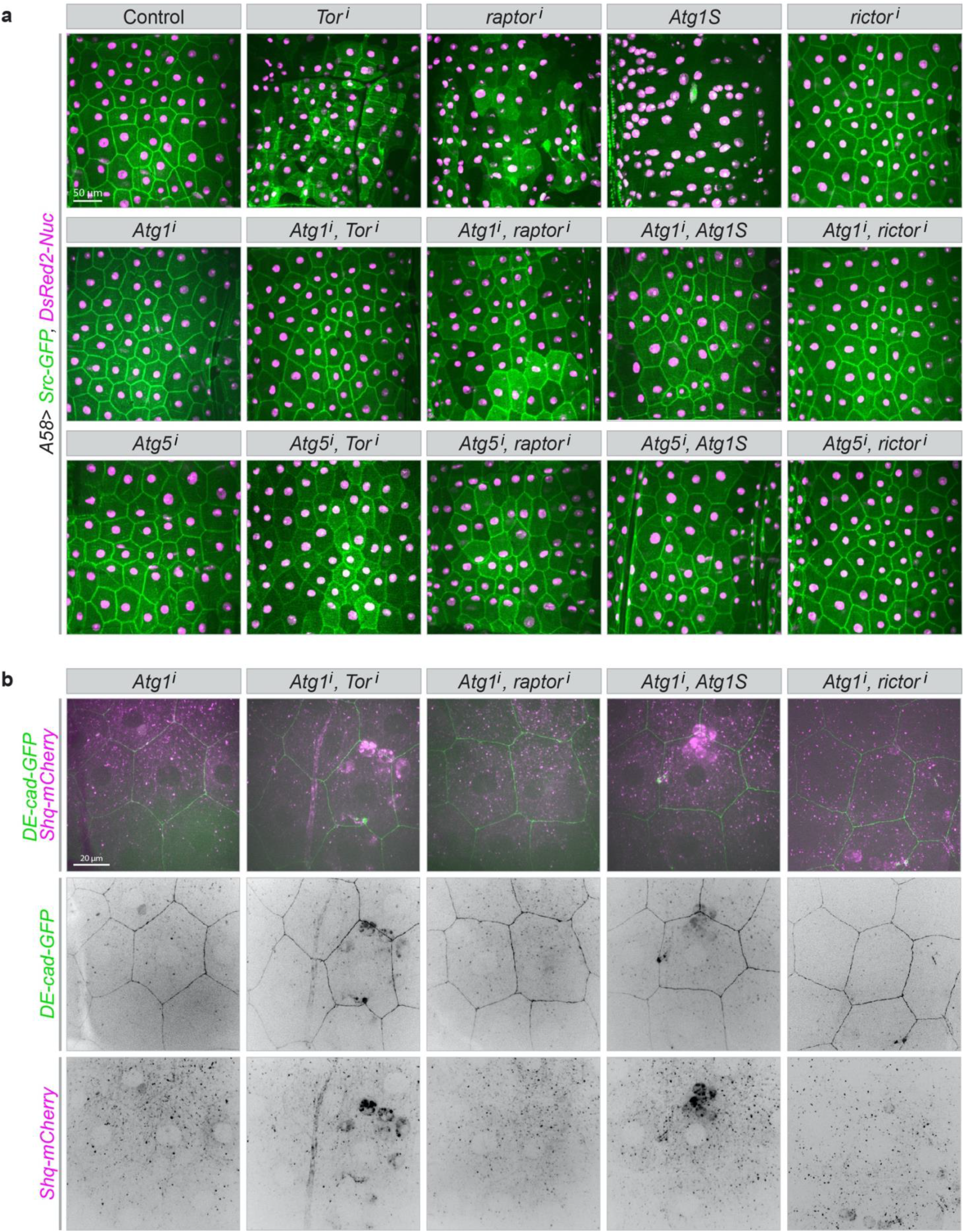
Epistasis of autophagy components in epidermal morphology. Larval epidermis expressing the indicated markers and overexpression or RNAi constructs. **a,** Inhibiting autophagy initiation (Atg1) or phagophore expansion (Atg5) in the epidermis suppresses the defects caused by loss of TOR signalling and uncontrolled autophagy. Lower magnification showing the entire width of larval segments A3 or A4 (compare to Fig. **7a**). **b,** Downregulating Atg1 suppresses the defects in E-cad and myosin distribution caused by down-regulation of TOR signalling (compare to **Fig. 4d** and **8a**). Images from **Supplementary Video 17**. **a,b,** n=10–3 larvae each genotype. **a,** 50 µm and **b,** 20 μm.

**Suppl. Fig. 9.**
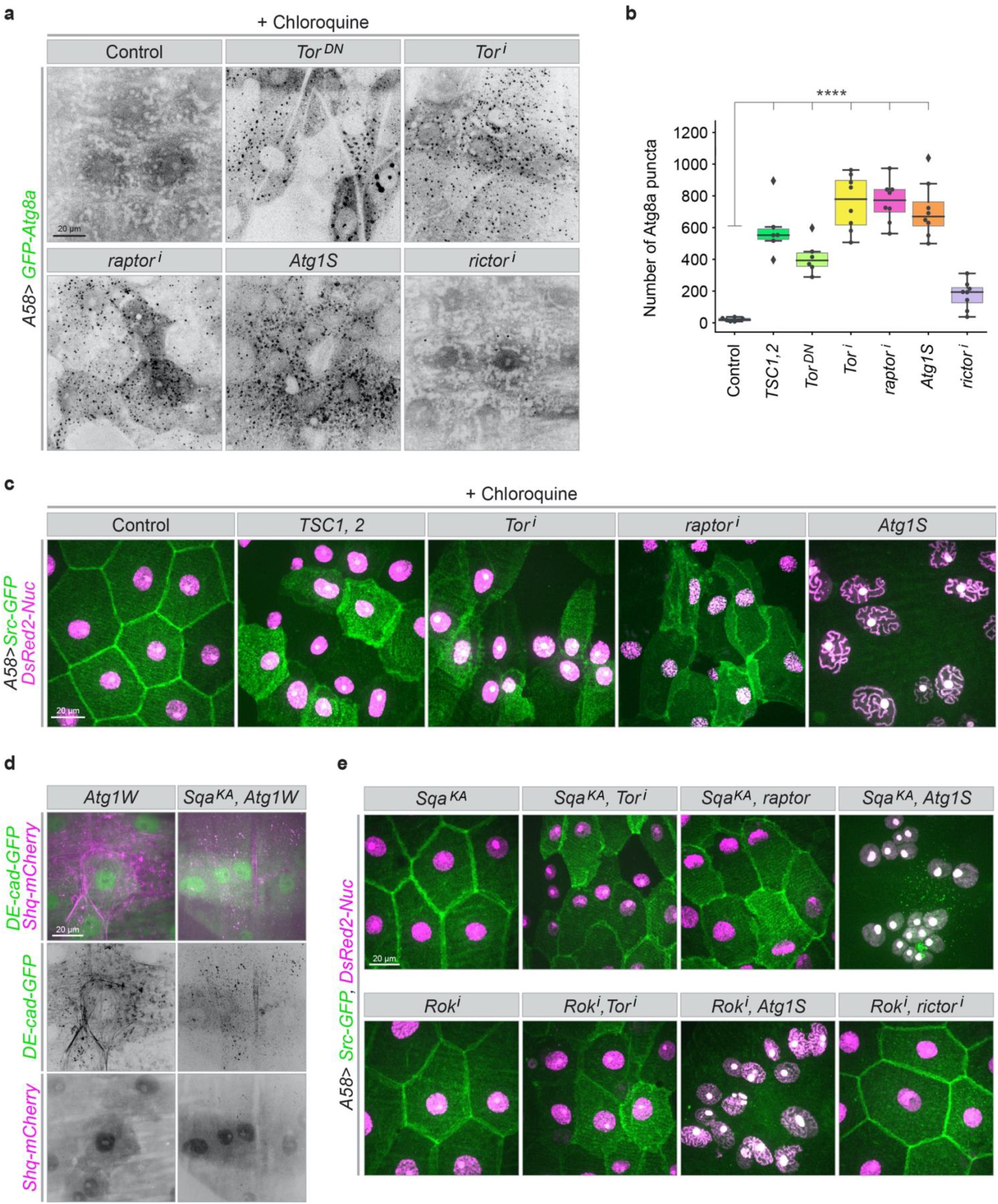
Role of autophagosome-lysosome fusion and myosin regulatory light chain activation on cellular defects. **a-c,** Epidermis expressing the indicated markers and constructs in larvae fed with chloroquine. **a,b,** Autophagosome formation in the presence of chloroquine is increased (compare to Fig. 3c). **b,** Quantification of autophagosomes in an epidermal area of 10000 µm^2^ in the indicated conditions, n=8 larvae each genotype. **c,** Blocking autophagosome maturation with chloroquine does not suppress the defects caused by uncontrolled autophagy. **d,** Blocking the myosin activator Sqa (Sqa^KA^), suppresses the formation of abnormal actin bundles but not the loss of lateral membranes (**e**). **c**, The effect of reduced TOR signalling cannot be suppressed by inhibiting Sqa or Rok. **a,c-e,** n=15–40 larvae each genotype. Scale bars: **a,c-e,** 20 μm.

**Suppl. Fig. 10.**
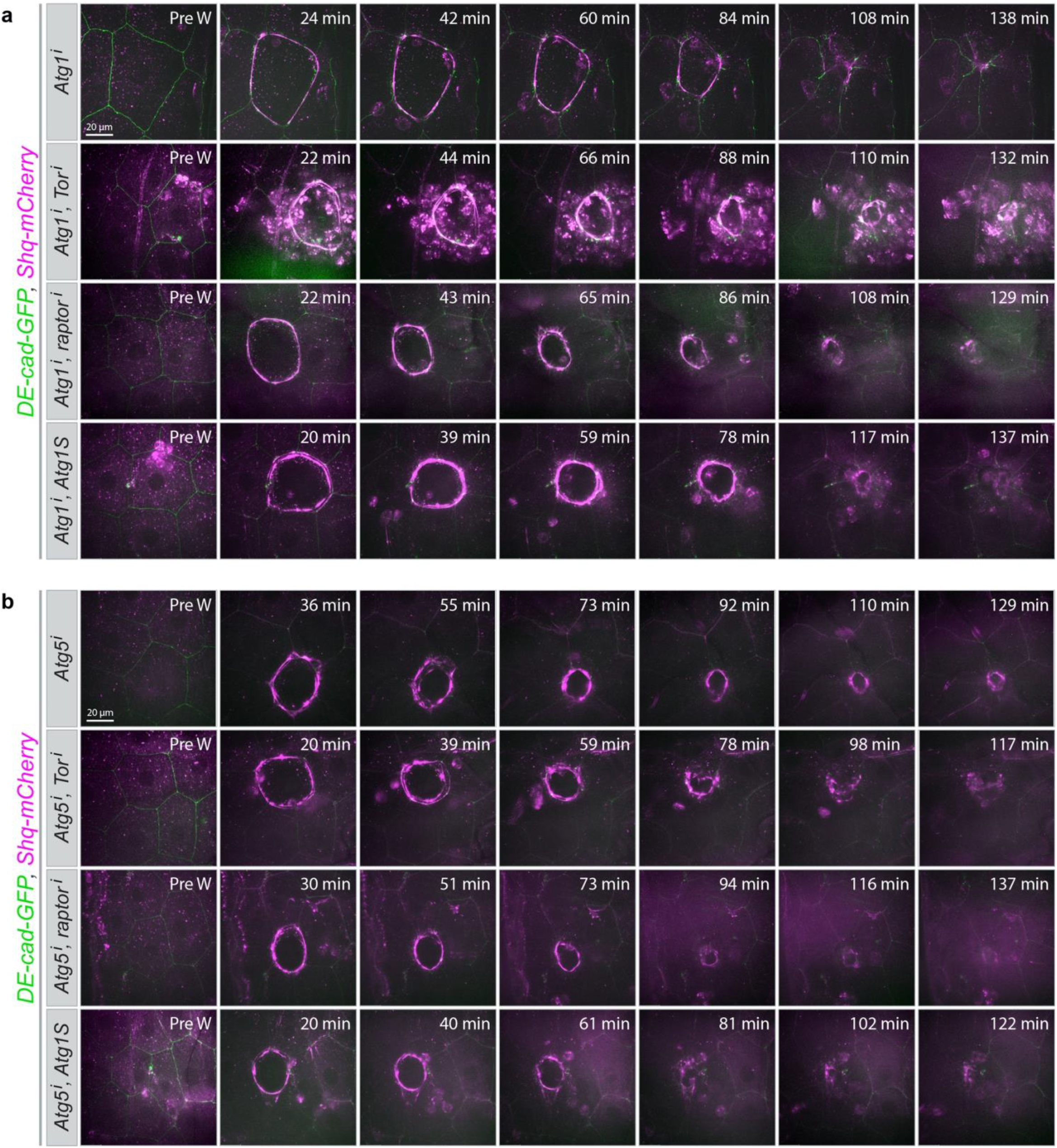
Role of autophagy in wound healing in TORC1 defective epidermis. **a,b,** Blocking autophagy alleviates wound healing defects caused by down-regulation of TOR signalling: timing of assembly, morphology and contraction of the actomyosin cable as well as wound closure are improved (compare to **Suppl.Fig. 4a**). **a,b,** n=5–20 larvae each genotype. Scale bars: **a,b,** 20 μm.

## Supplementary Videos

**Supplementary Video 1 | Autophagy during epidermal wound healing.** Live imaging of autophagic structures (marked with GFP-Atg8a) during wound closure in the epidermis of control (*A58>GFP-Atg8a*) larvae and after epidermal knockdown of Atg1, Atg5 and Atg6. Each frame shows a merged z-stack (see Materials and Methods). Scale bar 20 μm. Genotypes of all larvae are listed in **Table 1**.

**Supplementary Video 2 | Effect of autophagy on wound healing dynamics.** Live imaging of single-cell laser wounding in the dorsal epidermis of an L3 larva expressing Src-GFP (green, membrane marker) and DsRed2-Nuc (magenta, marker for nuclei) in the epidermis of control (*A58>Src- GFP,DsRed2-Nuc*) and after epidermal knockdown of Atg1, Atg5 and Atg6. Each frame shows a merged z-stack (see Materials and Methods). Scale bars: control, 11 μm and the others, 20 μm.

**Supplementary Video 3 | Effect of autophagy on actomyosin cable formation.** Live imaging of laser wounding in the epidermis of a larva expressing endogenously GFP-tagged E-cadherin (green) and Sqh-mCherry (mCherry-marked myosin regulatory light chain magenta) to visualize adherens junctions and actomyosin cables in control larva and after epidermal knockdown of Atg1 and Atg5. Each frame shows a merged z-stack (see Materials and Methods). Scale bar: 20 μm.

**Supplementary Video 4 | Syncytium formation during wound healing.** Live imaging of laser wounding in epithelia with clonally expressed free cytosolic GFP (magenta) under the *actin5c-Gal4* driver (*act>GFP,* Control). To visualize cell borders DE-cad-RFP (green) was ubiquitously expressed in all cells. **a,** Small clone: wound surrounded by two GFP expressing cells. **b,** Large clone: wound surrounded by more than two GFP expressing cells (in this case four cells are GFP positive). Each frame shows a merged z-stack (see Materials and Methods). Scale bar: 20 μm.

**Supplementary Video 5 | Testing GFP *de novo* synthesis in response to wounding or laser cut.** Live imaging of control larva (*act>GFP)* in which a clonally GFP-expressing cell (magenta) was wounded (**a**) or a central cell (GFP negative) was laser cut at the apical membrane without fatally wounding it (**b**). To visualize cell borders DE-cad-RFP (green) was expressed in all cells. Each frame shows a merged z-stack (see Materials and Methods). Scale bar: 20 μm.

**Supplementary Video 6 | Effect of supressing Atg1 on syncytium formation during wound healing.** Live imaging of laser wounding in epithelia with clonally expressed RNAi constructs for *Atg1* in GFP-positive clones (magenta, *act>GFP, Atg1^i^*). Cell borders are visualized by DE-cad-RFP (green). **a,** Small clone and **b,** large clone. Each frame shows a merged z-stack (see Materials and Methods). Scale bar: 20 μm.

**Supplementary Video 7 | Effect of supressing Atg5 on syncytium formation during wound healing.** Live imaging of laser wounding in epithelia with clonally expressed RNAi constructs for *Atg5* in GFP-positive clones (magenta, *act>GFP, Atg5^i^*). Cell borders are visualized by DE-cad-RFP (green). **a,** Small clone and **b,** large clone. Each frame shows a merged z-stack (see Materials and Methods). Scale bar: 20 μm.

**Supplementary Video 8 | Temporal effect of autophagy induction.** Live imaging after time-controlled overexpression of Atg1S together with Src-GFP (green, membrane marker) and DsRed2-Nuc (magenta, marker for nuclei) in the larval epidermis. Gene expression is induced 6 h before live imaging (see the schematic in Fig. 3g). Each frame shows a merged z-stack (see Materials and Methods). Scale bar: 20 μm.

**Supplementary Video 9 | Effect of uncontrolled autophagy on actomyosin cable formation during wound healing.** Live imaging of laser wounding in the epidermis of larva expressing endogenously DE-cad-GFP (green) and Sqh-mCherry (magenta) to visualize adherens junctions and actomyosin cables in control larva and after ectopic expression of Atg1W in the epidermis. Each frame shows a merged z-stack (see Materials and Methods). Scale bar: 20 μm.

**Supplementary Video 10 | Effect of Tor signalling and uncontrolled autophagy on actomyosin cable formation.** Live imaging of laser wounding in the epidermis of larva expressing endogenously DE-cad-GFP (green) and Sqh-mCherry (magenta) in control larva and after epidermal expression of Atg1S or RNAi constructs specific for Tor. Each frame shows a merged z-stack (see Materials and Methods). Scale bar: 20 μm. Scale bars: Tori 20 μm and Atg1S, 19 μm.

**Supplementary Video 11 | Effect of uncontrolled autophagy on wound healing.** Live imaging of laser wounding in control epidermis and after epidermal expression of Atg1S. Src-GFP (green) marks the plasma membrane and DsRed2-Nuc (magenta) marks the nuclei. Each frame shows a merged z-stack (see Materials and Methods). Scale bar: 20 μm.

**Supplementary Video 12 | Effect of uncontrolled autophagy on lateral membrane integrity.** Live imaging of FLIP experiment to test free cytoplasmic GFP motility within the epidermis of control larva (**a**) and after overexpression of Atg1W in epidermal cells (**b,c**). **a,b,c,** An area (magenta circle) was laser-illuminated for 3 min and recovery was imaged for 20 min after bleaching. Bleached area: **a,b**, 179 µm^2^; **c,** 1098 µm^2^. Each frame shows a merged z-stack (see Materials and Methods). Scale bars: control, 19 μm and the others, 20 μm.

**Supplementary Video 13 | Effect of uncontrolled autophagy on membranes.** Live imaging of FLIP experiment to visualise GFP motility within the control and Atg1W expressed epidermis. Z-section view of control and Atg1S overexpressing epidermis no leakage of GFP through the apical and basal membrane are seen. Each frame shows a merged z-stack (see Materials and Methods). Scale bar: 10 μm.

**Supplementary Video 14 | Uncontrolled autophagy and membrane leakage.** Live imaging of FLIP experiment to test GFP motility within the epidermis of two larvae expressing Atg1W (**a,b**). An area of 1098 µm^2^ (magenta circle) was laser-illuminated for 20 min and recovery was imaged for an additional 20 min after bleaching. Each frame shows a merged z-stack (see Materials and Methods). Scale bar: 19 μm.

**Supplementary Video 15 | Effect of uncontrolled autophagy on nuclei.** Live imaging of nuclei in epidermal cells expressing Atg1S and DsRed2-Nuc (black) as marker for nuclei. Each frame shows a single plane. Scale bar: 20 μm.

**Supplementary Video 16 | Atg1 acts via Atg5.** Live imaging of FLIP experiment shows that epidermal knockdown of Atg5 abolishes Atg1-induced syncytium formation. Each frame shows a merged z-stack (see Materials and Methods). Scale bar: 20 μm.

**Supplementary Video 17 | Block of Atg1 improves actomyosin cable and wound healing in uncontrolled autophagy.** Live imaging of laser wounding in the epidermis of control larvae expressing endogenously GFP-tagged DE-cad (green) and Sqh-mCherry (magenta) and after epidermal expression of an RNAi construct for Atg1 alone, or together with raptor RNAi or overexpression of Atg1S. Each frame shows a merged z-stack (see Materials and Methods). Scale bar: 19 μm.

**Supplementary Video 18 | Block of Atg5 improves actomyosin cable and wound healing in uncontrolled autophagy.** Live imaging of laser-wounding in the epidermis expressing endogenously GFP-tagged DE-cad (green) and Sqh-mCherry (magenta) in control larvae and after epidermal expression of an Atg5 RNAi construct alone, or together with Tor RNAi or overexpression of Atg1S. Each frame shows a merged z-stack (see Materials and Methods). Scale bar: 19 μm.

